# Systematic variation in marine dissolved organic matter stoichiometry and remineralization ratios as a function of lability

**DOI:** 10.1101/710178

**Authors:** Emily J. Zakem, Naomi M. Levine

## Abstract

Remineralization of organic matter by heterotrophic organisms regulates the biological sequestration of carbon, thereby mediating atmospheric CO_2_. While surface nutrient supply impacts the elemental ratios of primary production, stoichiometric control by remineralization remains unclear. Here we develop a mechanistic description of remineralization and its stoichiometry in a marine microbial ecosystem model. The model simulates the observed elemental plasticity of phytoplankton and the relatively constant, lower C:N of heterotrophic biomass. In addition, the model captures the observed decreases in DOC:DON and the C:N remineralization ratio with depth for more labile substrates, which are driven by a switch in the dominant source of labile DOM from phytoplankton to heterotrophic biomass. Only a model version with targeted remineralization of N-rich components is able to simulate the observed profiles of preferential remineralization of DON relative to DOC and the elevated C:N of bulk DOM. The model suggests that more labile substrates are associated with C-limited heterotrophic growth and not with preferential remineralization, while more recalcitrant substrates are associated with growth limited by processing rates and with preferential remineralization. The resulting patterns of variable remineralization stoichiometry mediate the extent to which a proportional increase in carbon production resulting from changes in phytoplankton stoichiometry can increase the efficiency of the biological pump. Results emphasize the importance of understanding the physiology of both phytoplankton and heterotrophs for anticipating changes in biologically driven ocean carbon storage.

## 1 Introduction

Primary production converts inorganic carbon, nitrogen, and other nutrients into biomass. The return reaction – the remineralization of organic matter back into its inorganic constituents by heterotrophic organisms – fuels this production by resupplying inorganic nutrients, thereby closing the most significant loop of biogeochemical cycling. In the ocean, nitrogen, phosphorus, or iron usually limits primary production (Moore et al. 2013), and so the rate at which phytoplankton fix carbon relative to the rate at which they utilize other elements controls the amount of organic carbon produced. In contrast, carbon is thought to limit heterotrophic activity, since heterotrophs generally excrete N and P without a shift in oxidation state, while oxidizing organic carbon to CO_2_ for energy. The C limitation of heterotrophs appears to be at odds with the large pool of dissolved organic carbon stored in the ocean (about 700 Pg C) (Hansell 2013).

Observations and modeling efforts have demonstrated global-scale patterns in the stoichiometry of plankton biomass, particulate and dissolved organic matter, and exported (sinking) organic matter in the ocean that vary significantly from the average “Redfield ratio” (C:N:P = 106:16:1) (e.g. Schneider et al. 2003; Aminot and Kérouel 2004; Martiny et al. 2013a,b; Teng et al. 2014; Letscher and Moore 2015; Devries and Deutsch 2014; Talmy et al. 2016; Kwiatkowski et al. 2018). In general, where primary production is more N and/or P limited, such as in oligotrophic subtropical regions, phytoplankton biomass is enhanced in C (Moreno and Martiny 2018). This allows for a higher flux of C to the deep ocean for a given amount of N- or P-based production (Teng et al. 2014; Devries and Deutsch 2014).

This flexibility has been hypothesized as an important negative feedback to changes in global climate. Galbraith and Martiny (2015) demonstrated a heightened sensitivity of ocean C storage and thus atmospheric CO_2_ to changes in ocean circulation when variable phytoplankton stoichiometry is considered. In addition, Moore et al. (2018) predicted large decreases in primary productivity at low latitudes due to global warming as a result of decreased surface nutrient supply. Together, these studies suggest that even though C production may decrease in a warmer world, the rate of decrease may be slower than that of N and P, therefore maintaining a relatively stable rate of C export to the deep ocean.

The impact of heterotrophic activity on this feedback remains unclear. Heterotrophs drive the rates, locations, and stoichiometry of the remineralization of organic matter. Thus, variations in these processes have the potential to impact biologically driven carbon storage and enhance or dampen the climate feedbacks anticipated from phytoplankton plasticity (Kwon et al. 2009). While C:N ratios of phytoplankton biomass vary widely with nutrient supply (Goldman et al. 1979; Martiny et al. 2013a; Galbraith and Martiny 2015; Talmy et al. 2016), heterotrophic bacteria seem subject to a different set of controls, since marine heterotrophic biomass on average has a lower and a more consistent C:N of 5±1 (Zimmerman et al. 2014). Even though bacterial carbon demand is high relative to N and P due to low growth efficiencies, the preferential remineralization of organic N and P relative to C by heterotrophs has often been inferred from observations (Schneider et al. 2003; Coles and Hood 2007; Landolfi et al. 2008; Burkhardt et al. 2014; Monteiro and Follows 2012; Letscher and Moore 2015). Here we define preferential remineralization as the difference between an elemental ratio of a specified organic pool and the ratio of its consumption. Thus, preferential remineralization of N relative to C means that the C:N of consumption is lower than the C:N of the consumed pool (i.e. ΔDOC:ΔDON < DOC:DON).

Letscher and Moore (2015) identified two changes in the stoichiometry of dissolved organic matter (DOM) with depth from the surface into the thermocline (Fig. 1), which reflect the activity of heterotrophic and other non-photosynthetic organisms. First, the C:N of the entire DOM bulk pool increases with depth, which Letscher and Moore (2015) concluded reflects an increasing fraction of C-rich recalcitrant substrates with depth in agreement with previous analysis (Aminot and Kérouel 2004). Second, the C:N of the “non-recalcitrant,” more quickly degraded portion of the DOM pool decreases with depth. Letscher and Moore (2015) then analyzed these patterns to estimate the non-recalcitrant DOM remineralization ratios and the degree of preferential remineralization of one element over another. They found that the C:N of remineralization decreased with depth, from above to below Redfieldian values. However, they only found evidence for preferential remineralization of N relative to C. DOC was not preferentially remineralized, despite a high C:N remineralization ratio at the surface, because the ratio matched the high C:N of the DOM itself. In contrast, the degree of preferential N remineralization relative to C 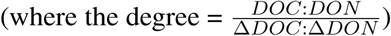 generally increased with depth.

**Figure 1:**
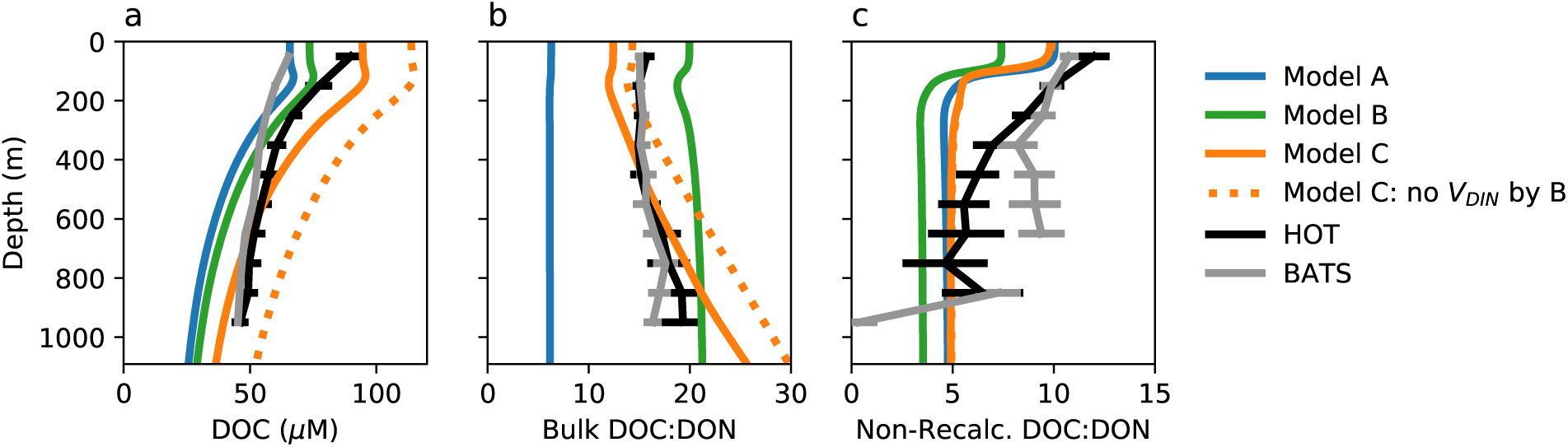
Observed and modeled DOC concentration and DOC:DON (mol/mol) patterns. (a) The mean and s.d. of DOC, and (b, c) the mean and s.d. of DOC:DON of eight annual profiles assembled by Letscher and Moore (2015) are shown for two ocean time series stations: HOT (the Hawaii Ocean Time-series) and BATS (the Bermuda Atlantic Time-series Station). As in Letscher and Moore (2015), non-recalcitrant DOM was estimated by subtracting out the lowest concentration of DOC or DON. The modeled non-recalcitrant DOM consists of the sum of the DOM classes that do not accumulate throughout the water column (the 15 fastest out of the 25 classes).

Why does the C:N of remineralization change with depth, and what controls the degree of preferential remineralization? We currently lack a mechanistic understanding of how heterotrophic activity results in the patterns identified by Letscher and Moore (2015), and so cannot predict how remineralization dynamics may change in the future. Understanding the patterns of DOM stoichiometry and remineralization requires resolving the stoichiometry of the consumption of organic substrates by heterotrophic prokaryotes. A suitably mechanistic description must also consider the heterogeneity of organic substrates, as well as reconcile the seeming paradox between the preferential remineralization of N and P and the carbon-limited growth of heterotrophs.

Galbraith and Martiny (2015) highlight the need for biogeochemical models to mechanistically resolve flexible stoichiometry in order to better understand and systematically predict changes in carbon cycling and climate feedbacks. Here, we work towards this goal by developing a dynamic microbial ecosystem model that allows for flexible elemental ratios of consumption, biomass synthesis, and respiration, as well as a spectrum of DOM pools with wide-ranging values of lability. Since we do not here focus on particulate organic matter (POM) stoichiometry, we resolve POM simplistically in order to isolate the direct controls on DOM. We investigate how microbial heterotrophs impact the stoichiometry of marine DOM, specifically focusing on the observed patterns of DOC:DON (Fig. 1), by studying the emergent ecosystem structure in simulations of a vertical marine water column.

## 2 Methods

Here, we first describe the metabolic model developed to resolve both phytoplankton and heterotrophic microbial activity. As the goal is to understand the distinctions in biomass C:N between phytoplankton and heterotrophs and their effects, we assume that both phytoplankton and heterotrophs have identical elemental ratios of ‘core’ biomass, and differ only in their means of energy acquisition. Second, we describe the treatment of DOM, for which discrete pools are resolved along a lability continuum. We here consider only the portion of DOM that originates from microbial biomass, though black carbon, terrestrial sources, and other recalcitrant compounds additionally contribute to the total pool in the ocean. Third, we describe the different controls governing the stoichiometry of uptake, including the ability of heterotrophs to take up inorganic nutrients. In this third section, we also describe three model versions that incorporate plausible controls on DOM stoichiometry (Models A, B, and C). Lastly, we explain the ecosystem configuration in an idealized stratified marine water column model.

### 2.1 Microbial metabolic model

We develop an internal stores (‘Droop,” or “quota”) model of heterotrophic and photoautotrophic microbial populations that resolves dynamic fluxes of C, N, and P through biomass synthesis, respiration, and excretion of waste products (Fig. 2a; Caperon 1968; Droop 1968; Caperon and Meyer 1972; Droop 1973). The quota model decouples the uptake of substrate from growth: uptake is a function of external (environmental) concentrations and stoichiometry, while growth is a function of internal (cellular) elemental quotas. This framework predicts internal accumulations of non-limiting, “excess” nutrients without specifying the physiological reasons for the accumulation. Quota models have been used to predict extra carbon in phytoplankton biomass in oligotrophic areas (e.g. Talmy et al. 2016; Kwiatkowski et al. 2018), but, to our knowledge, have not been used in conjunction with models of marine heterotrophic growth. The model presented here follows that of Thingstad (1987) with three main extensions: i) the excretion of N and P as well as C as waste products, ii) the ability of heterotrophs to take up inorganic N and P to supplement biomass synthesis, and iii) the description of photoautotrophic growth and excretion using the same set of equations.

**Figure 2:**
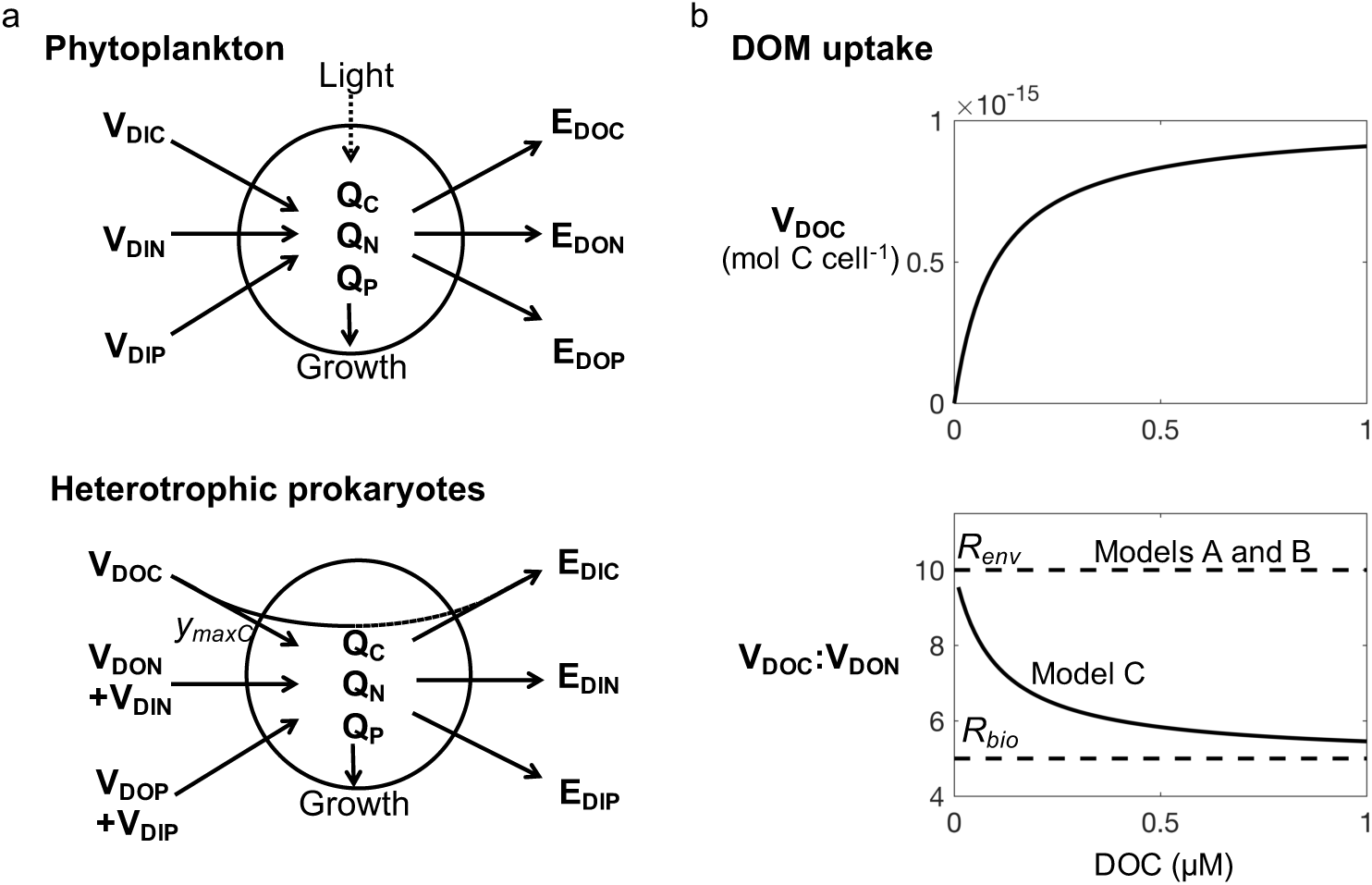
Metabolic model for photoautotrophic and heterotrophic metabolism. (a) Schematic of the internal stores model of cell metabolism for phytoplankton and heterotrophic prokaryotic populations, with uptake rate *V*, excretion rate *E*, and cell quota *Q*. For heterotrophs, a mandatory maximum C yield (*y*_*maxC*_) represents the required utilization of organic C for energy (respiration). For light-harvesting phytoplankton, *y*_*maxC*_ = 0, and an additional light limitation on growth is applied. (b) Illustration of the model parameterizations for DOM uptake and the C:N of uptake for the case where the environmental ratio *R*_*env*_ ([DOC]:[DON]) is 10. DOC uptake is illustrated for specific uptake rate 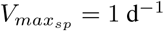.

Following the quota model formulation, we describe the dynamics of microbial population *i* as

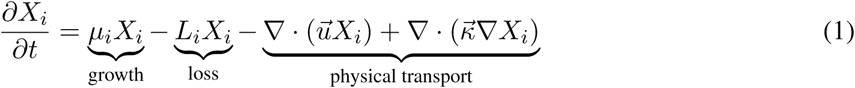

where *X*_*i*_ (cells L^−1^) is the cell abundance of each population that changes with growth rate *μ*_*i*_ (t^−1^), loss rate *L*_*i*_ (t^−1^), and physical transport. We then account for the biomass *B*_*i,j*_ (mol L^−1^) associated with each element *j* (where *j* is C, N, or P), as

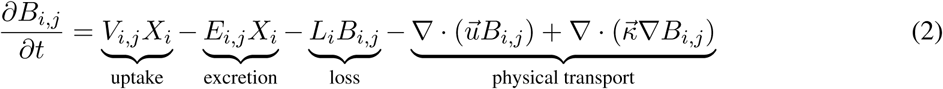

where *V*_*i,j*_ (mol cell^−1^ t^−1^) is the total uptake of each element, and *E*_*i,j*_ (mol cell^−1^ t^−1^) is excretion. Biomass and abundance are related by the cell quota of each element *Q* (mol cell^−1^) as *B*_*i,j*_ = *X*_*i*_*Q*_*i,j*_. The limiting cell quota controls the growth rate relative to the minimum cell quota 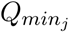 (mol cell^−1^) of element *j* as:

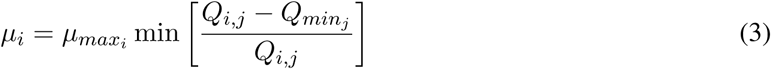

Here, we assume identical minimum quotas for each element for all populations of phytoplankton and heterotrophs. Assigned ratios of the minimum quotas for each element (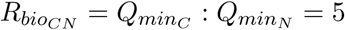 and 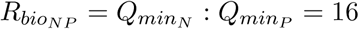) set the ‘core’ biomass stoichiometry.

Total uptake *V*_*i,j*_ is the summed uptake of one or more substrates containing element *j* (such as DON and 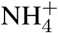 for total N uptake). For each substrate *S* (mol L^−1^), we describe uptake *V*_*S*_ (mol cell^−1^ t^−1^) with a Michaelis-Menten form as:

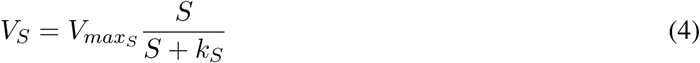

where 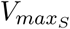 (mol cell^−1^ t^−1^) is a maximum cellular uptake rate and *k*_*S*_ (mol L^−1^) is a half-saturation concentration. Thus 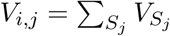. Phytoplankton take up elements only in inorganic form, though in the future this model could be extended to consider organic assimilation by phytoplankton. Heterotrophs take up elements preferentially in organic form, though we also investigate their ability to take up inorganic N and P to supplement elemental demands for biomass synthesis.

Following Thingstad (1987), we calculate the excretion *E*_*i,j*_ (mol cell^−1^ t^−1^) of each element out of the cell as a function of two terms: the excess quota of each element and the amount of uptake allocated for respiration by the cell, as:

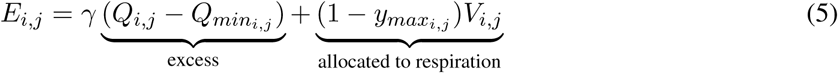

where *γ* (t^−1^) regulates the rate of excretion of excess element, and where *y*_*max*_ is a maximum yield of each element (mol biomass per mol uptake) that corresponds to the minimum portion of substrate taken up by the cell that must be used for respiration for energy. The parameter *γ* thus controls the degree of ‘leakiness’ of the cells. Here, we resolve heterotrophic excretion of nutrients in inorganic form, though the model could be extended to investigate succession dynamics by resolving the excretion of organic intermediates by different populations.

For heterotrophs, we assume that *y*_*max*_ = 1 for N and P, and that *y*_*max*_ < 1 for C, since organic carbon serves as the electron donor for respiration. For the water column model, we assign 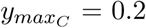 in accordance with an estimate of the portion of organic carbon consumed that must be respired for energy to fuel biomass synthesis using published marine values (Robinson and Williams 2005) (Table 1). For phytoplankton, we assume all energy is acquired from photons and thus that *y*_*max*_ = 1 for all elements (i.e., we resolve net primary production). Phytoplankton growth is additionally limited by light availability. An analysis of model sensitivity to 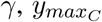, and other parameters shows that the resulting qualitative patterns are consistent across a plausible range of parameter values. (See Supporting Text S2 for all equations, parameter values, and sensitivity results.)

**Table 1:** Metabolic parameters for heterotrophs in the water column model. (See Supporting Information for complete list of model parameters and variations in values.)

| Parameter | Symbol | Value | Units |
| --- | --- | --- | --- |
| C:N ratio of minimum quotas ($Q_{minC} : Q_{minN}$) | $R_{bio_{CN}}$ | $5^{\dagger}$ | mol/mol |
| N:P ratio of minimum quotas ($Q_{minN} : Q_{minP}$) | $R_{bio_{NP}}$ | $16^{\dagger}$ | mol/mol |
| Minimum N quota | $Q_{minN}$ | $0.2^{\ddagger*}$ | fmol N cell<sup>-1</sup> |
| Maximum carbon yield | $y_{maxC}$ | $0.2^{\S*}$ | unitless |
| Maximum specific uptake rate | $V_{max_{sp}}$ | $0.01 - 100$ | d<sup>-1</sup> |
| Half-saturation constant, DOC | $k_{DOC}$ | $0.5V_{max_{sp}}^*$ | μM C |
| Maximum growth rate | $\mu_{max}$ | $\max(1, V_{max_{sp}})^*$ | d<sup>-1</sup> |
| Regulation rate of excretion | $\gamma$ | $0.5^*$ | d<sup>-1</sup> |
| Quadratic mortality rate | $m_q$ | $1^*$ | μM N<sup>-1</sup> d<sup>-1</sup> |

<sup>†</sup>Anderson (1995)
<sup>‡</sup>Bratbak and Dundas (1984) with a cell diameter of 0.5 μm (Sherr and Sherr 2000)
<sup>§</sup>Robinson and Williams (2005)
\*Variations: Figs. S6, S9–S11 (Table 2)

### 2.2 A spectrum of organic matter lability

Despite its immense complexity, DOM is often conceptually organized along an axis of ‘lability,’ where the rate of microbial degradation of a substrate (or set of substrates) correlates with the degree of its lability (Hansell 2013). For less labile, ‘recalcitrant,’ substrates, growth may be limited due to a low energy content and/or a high cost of accessibility (LaRowe and Van Cappellen 2011). Here, we define lability *a priori* with generalized uptake kinetic parameters. In reality, lability is an emergent property that is affected by kinetic constants of enzymes and by the phenotypic state of the heterotrophs in terms of the amount of metabolic machinery devoted to the consumption of a given organic substrate. We do not resolve these processes explicitly, but instead prescribe lability as an aggregated characteristic that reflects their effects. In this way, we consider lability as a property not solely of the DOM pool or the heterotrophic population that consumes it, but as an ‘ecosystem property’ that reflects both. This is similar to a typical biogeochemical model that resolves slow, medium, and fast remineralization timescales of DOM (and POM), and to a “multi-G” model that resolves a continuum of remineralizgeation rates (Middelburg 1989; Aumont et al. 2017).

We discretize DOM into 25 classes meant to represent a diversity of organic compounds that are consumed at similar rates. Each class is associated with a specific maximum rate of uptake 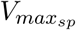 that varies from 0.01 d^−1^ to 100 d^−1^ (Fig. 3). 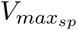 represents the integrated impact of multiple cellular processes. Since we are not investigating trait-based competition, for simplicity, we assume each DOM class is associated with the same uptake affinity 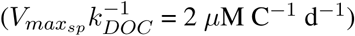 Each class consists of three distinct pools of DOC, DON, and DOP that are resolved independently as state variables. The choice of 25 classes is sufficient to examine the relationship between the rate and the stoichiometry of remineralization while remaining computationally tractable in one dimension.

**Figure 3:**
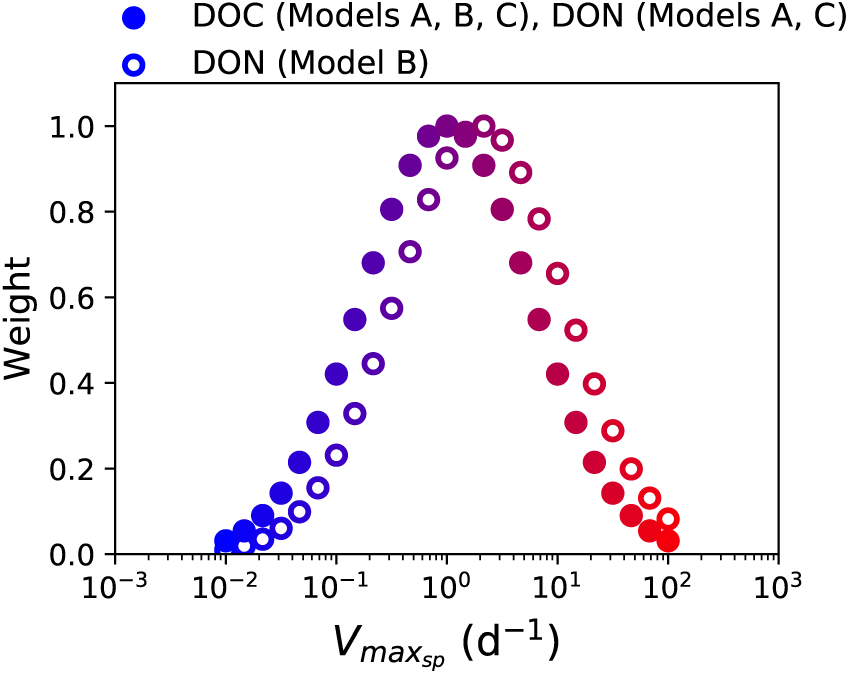
DOM lability and its distribution in the ecosystem model. Lability is defined by the specific maximum uptake rate 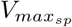 of 25 discrete classes of DOM. The distribution represents the partitioning of the production of DOM among the 25 pools. In Model B, an intrinsically higher lability of DON than DOC is represented with a higher mean distribution. This results in a higher initial C:N of the less labile substrates, and a lower initial C:N of the more labile substrates, in proportion to the vertical differences between the solid and open circles. The illustrated color correlates with 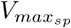 for ease of interpretation of later illustrations (Figs. 4, 5, and 6).

When DOM is produced in the model, it is partitioned into each of these 25 classes following an assumed distribution. Though the dynamics of organic matter degradation by a diverse heterotrophic community are complex, theory and measurements suggest that remineralization rates are lognormally distributed as a result of multiplicative stochastic processes (Rothman and Forney 2008; Forney and Rothman 2012; Rothman 2014; Middelburg 1989; Follett et al. 2014; Mostovaya et al. 2017). Such stochastic processes include the likelihood that the enzyme required to degrade a specific substrate is co-located with that substrate (Arnosti 2011), variation in the abundances of heterotrophic populations with diverse enzymatic capabilities, differences in the energetic content of substrates, differences in the accessibility or difficulty of degradation of a particular compound (perhaps requiring a more expensive enzyme than another), the cost of building enzymes for hydrolysis and transport, the encounter rates between heterogeneous organic molecules and biomass, etc. (Forney and Rothman 2012; Rothman 2014; Cael et al. 2018).

Here, we assume that the aggregated impact of these stochastic processes governing remineralization yields a lognormal distribution of the production of the 25 DOM classes. Thus, the majority of DOM produced is associated with a mean rate while much smaller amounts are associated with the fastest and slowest rates (Fig. 3). This assumed distribution aims to capture the predicted end-state of remineralization dynamics, and so we do not explicitly resolve many underlying processes such as the transformations of DOM between the classes. The lognormal distribution accounts for such transformations implicitly. Because of the lognormal scaling, this distribution is not directly proportional to the production timescales of the common DOM categories of “‘labile,” “semi-labile,” “semi-refractory,” “refractory,” and “ultra-refractory” as listed in Hansell (2013): rather, the majority of the classes fall into the “labile” category since the mean 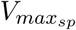 of 1 d^−1^ corresponds to relatively rapid consumption, while fewer classes are “refractory.” Qualitatively, the model solutions are independent of the form of the distribution (see Fig. S7 for the solutions with an even distribution, for example), but a lognormal distribution results in an improved match to observed DOC concentrations.

### 2.3 Stoichiometry of uptake

#### 2.3.1 Inorganic uptake

We parameterize the uptake of inorganic N and P following previous approaches used for phytoplankton (Follows et al. 2007; Ward et al. 2013). We use empirically based estimates for the maximum uptake rate and half-saturation concentration for dissolved inorganic nitrogen (DIN) (Litchman et al. 2007), and we assume that DIP uptake parameters relate stoichiometrically as the core biomass ratio 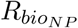 (Follows et al. 2007; Ward et al. 2013). The model solutions are not sensitive to the particular ratio of inorganic nutrient uptake parameters because phytoplankton deplete inorganic nutrients whenever light energy allows, and because inorganic nutrients do not impact the behavior of heterotrophs in the model. We then relate the maximum C fixation rate to that of DIN (and DIP) with 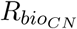, representing the ability of phytoplankton to assimilate DIC to proportionally match the maximum rate of DIN (and DIP) uptake (Supporting Text S2). This allows for excess C fixation, and subsequent DOC excretion for “leaky” phytoplankton cells, when DIN (or DIP) limits growth.

We allow heterotrophs to take up DIN (or DIP) when the uptake ratio of organic C:N (or C:P) exceeds 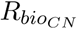 (or 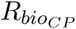), using the same parameters for inorganic uptake as for phytoplankton (Supporting Text S2). To test for competition between heterotrophs and phytoplankton for inorganic nutrients, we also run a comparative model in which heterotrophs do not take up DIN or DIP.

#### 2.3.2 Organic uptake by heterotrophs

Since DOM is composed of a heterogeneous mix of organic molecules, many microscale processes govern its uptake and modification by the heterotrophic community. Given this complexity, we consider that two extremes should govern the stoichiometry of uptake: regulation by the stoichiometry of the available substrate, and regulation by a biologically preferred ratio set by metabolic machinery. This corresponds to a dichotomy in marine microorganism strategy of “you are what you eat” vs. “you eat what you are” (Moreno and Martiny 2018).

If heterotrophic prokaryotes ‘eat what they are,’ they may utilize the N-rich components of organic compounds that generally have C:N ratios more similar to their requirement for biomass synthesis than to the pool of DOM, leaving excess DOC behind. Alternatively, they may have adapted to take into account the additional organic C required for respiration, and so take up C:N at a ratio higher than that of biomass synthesis. Such preferential consumption of either N or C is plausible since the uptake of organic substrate by prokaryotes involves the extracellular hydrolysis of organic molecules. Though we lack a comprehensive understanding of the stoichiometry of DOM uptake, regulations of uptake stoichiometry have been observed in experiments with E. Coli and natural marine assemblages (You et al. 2013; Burkhardt et al. 2014).

Heterogeneity in the ‘quality’ of organic substrate should also impact uptake stoichiometry. The global patterns revealed by Letscher and Moore (2015) suggest that a different stoichiometric control may govern the consumption of non-recalcitrant vs. recalcitrant substrates. If C-rich compounds are on average less labile than N-rich compounds upon formation (e.g. cell walls vs. amino acids), then the C:N of DOM may be enhanced simply due to this supply stoichiometry. If heterotrophs preferentially target N-rich compounds or fragments of compounds, then the C:N of uptake will be lower than that of the DOM itself, constituting preferential remineralization.

The above controls are not mutually exclusive and, most likely, co-occur to produce the observed patterns. However, to robustly test the proposed mechanisms, we construct three ‘end-member’ models for the uptake of DOC and DON: 1. (Model A) uptake stoichiometry matches that of the available organic substrate, 2. (Model B) uptake stoichiometry matches that of the available organic substrate, with DON on average more labile than DOC, and 3. (Model C) uptake stoichiometry matches that of the available organic substrate when its concentration limits growth, but is regulated biologically to match a preferred uptake ratio when growth is limited by the consumption rate of the substrate.

We do not consider two other possible models because they result in unrealistic solutions. First, the stoichiometry of uptake could match the biologically preferred ratio *R*_*bio*_ at all concentrations. However, this results in the accumulation of labile DOC when DOC:DON *> R*_*bio*_, which is not consistent with observations. Second, organisms may have adapted to take into account the extra C required to satisfy energetic demand via respiration, such that uptake stoichiometry would be 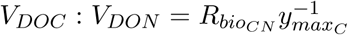. With typical values of 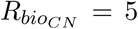 and 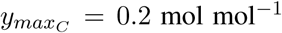, this suggests that *V*_*DOC*_ : *V*_*DON*_ ≈ 25, which results in preferential remineralization of C rather than of N and P. This is the opposite of what is observed (Letscher and Moore 2015), and thus is not a suitable general parameterization for all DOM classes. However, this preferential uptake of C may indeed characterize the uptake of labile carbon substrates such as glucose. As described below, Model C accounts for this latter case by assuming that uptake stoichiometry increasingly matches availability as substrates are depleted, allowing for high C:N of uptake for labile substrates.

For models A, B, and C, we describe the uptake of each class of DOC with a saturating (Michaelis-Menten) form, in terms of a half-saturation constant *k*_*DOC*_ and a maximum uptake rate 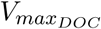 (mol C cell^−1^ d^−1^). This cellular maximum uptake rate can be decomposed as:

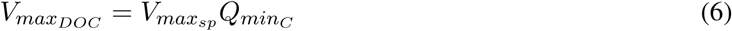

For model versions A, B, and C, the uptake parameters for DON (and DOP, analogously) are defined below (Fig. 2b):

##### Model A: Uptake stoichiometry matches the stoichiometry of available DOM

For the first end-member, the parameters for uptake of DON are related to those of DOC as:

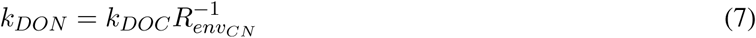

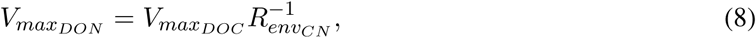

where 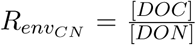. With this parameterization, the ratio of uptake matches the ratio of the ambient DOM pool. The stoichiometry of uptake therefore adjusts dynamically to changes in the DOM stoichiometry (driven by supply stoichiometry), and the consumption of DOM by heterotrophs has no influence on the stoichiometry of DOM (Fig. 2b).

##### Model B: Uptake stoichiometry reflects DOC:DON lability

In this end-member, we assume that DON has an inherent higher lability than DOC. This model represents differences in the C:N of DOM at production, rather than at consumption, as a function of lability. We represent this higher lability simply by assigning a higher lognormal mean of the distribution of DON compared to DOC (Fig. 3), i.e., we impose this difference in lability during DON formation. Thus, as DOM is supplied from biomass mortality, more DON is partitioned into the classes associated with faster uptake kinetics. As a consequence, the C:N of more recalcitrant substrates is initially higher, and the C:N of more labile substrates is initially lower (Fig. 3). To isolate the impact of this, we assume that uptake matches DOM stoichiometry as in Model A, and so uptake itself does not impact the DOM stoichiometry.

##### Model C: Uptake stoichiometry of accumulated substrates matches biological demand

For this end-member, uptake stoichiometry reflects a biologically regulated ratio of demand. Here, we assume that this biologically regulated ratio is 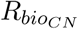, and we relate the uptake kinetic parameters with this ratio as:

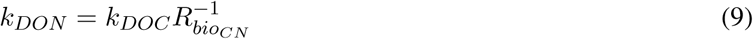

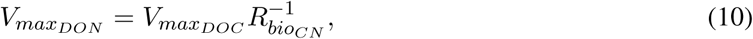

Unlike Models A and B, this parameterization results in uptake stoichiometry that differs from ambient DOM stoichiometry, and varies as a function of DOM concentrations. Specifically, uptake stoichiometry matches that of external organic matter stoichiometry when concentrations are low, and converges to 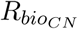 when organic matter concentrations are high relative to half-saturation concentrations (Fig. 2b). In the water column, if the accumulation of organic substrates inversely correlates with their lability, then this parameterization allows for labile DOC uptake to reach high C:N uptake ratios, while recalcitrant DOC is discriminated against relative to DON consumption.

### 2.4 Water column configuration

We simulate the ecosystem dynamics of a vertical marine water column, resolving C, N, and P cycling independently. The model represents the three species of DIN (NH_4_, NO_2_, and NO_3_), DIP (PO_4_), and 25 pools of DOM which is traced as DOC, DON, and DOP. Because we focus on DOM heterogeneity, we resolve only one class of POM sinking at a constant rate and consumed by one POM-associated heterotrophic population. Similarly, for simplification, we resolve only one phytoplankton functional type that can assimilate NH_4_, NO_2_, and NO_3_ as in Zakem et al. (2018) (see Supporting Text S2 for detail). We resolve heterotrophic biomass as ‘populations’ that are each sustained by one or more of the 25 DOM classes. All heterotrophs are allowed to assimilate NH_4_, NO_2_, and NO_3_ and two explicit populations of nitrifying microorganisms using the formulation in Zakem et al. (2018) are also included. The speciation of DIN and the elemental stoichiometry of the nitrifiers do not impact solutions (model results are identical if just one pool of DIN is resolved). Following the Droop model formulation, each population is represented by four state variables: C, N, and P biomass (*B*_*C*_, *B*_*N*_, and *B*_*P*_), and cell density *X* (Eqns. 1 and 2).

The 25 pools of biomass are not meant to strictly represent actual heterotrophic groups, and so the number of co-existing populations is not strictly related to the diversity of the community, but it does crudely represent the amount of biomass sustained on “labile” vs. “recalcitrant” DOM. We explore the sensitivity of the model results to different formulations of the subsets of DOM that sustain these biomasses. We resolve a ‘specialist’-like configuration in which the 25 biomass pools are each sustained by one class of DOM (as illustrated in the main text), a ‘generalist’-like configuration in which one biomass population consumes all 25 classes, and a combined configuration in which biomasses that are able to consume less labile substrates may also consume the more labile substrates, at the expensive of a lower maximum growth rate (Figs. S12 and S13).

Losses to grazing and viral lysis are represented with a quadratic mortality term as *L*_*i,j*_ = *m*_*q*_*B*_*i,j*_, where *m*_*q*_ is a quadratic mortality parameter, identical for all populations. Both POM and DOM are produced from the losses of all biomasses, and partitioned according to parameter *f*_*mort*_. DOM is additionally produced from POM via a ‘sloppy grazing’ that represents the amount of organic molecules that escape uptake following extracellular hydrolysis, for which a specified amount of DOM is transformed from POM per mole POM consumed, according to parameter *α*. DOM production is partitioned among the 25 pools following the distribution in Fig. 3.

To give the water column its physical structure, a mixed layer is simulated with a vertical diffusion coefficient, which attenuates with depth to a minimum that represents eddy-driven vertical mixing in the thermocline (Supporting Text S2, Table S2). Light energy decreases with depth according to the attenuation coefficient for water. All populations and nutrients are vertically mixed, and POM is additionally advected according to its sinking rate. The model includes a total of 197 state variables. The model is run to a steady state (*>* 3000 years), defined as < 1% change in total DOC per year.

We explored the sensitivity of ecosystem model to confirm that plausible variations in the parameter values (listed in Table 1 and Table S2) do not affect solutions qualitatively (Figs. S3–S11). Results were consistent across all parameters. In addition, results were also robust across the different ecosystem configurations of DOM consumption by heterotrophic biomass pools as long as two conditions are met: that some pools of organic matter accumulate throughout the water column, and that some pools are depleted to low concentrations. We found that all configurations met these conditions, if populations consuming multiple substrates were penalized, and thus gave qualitatively similar solutions except for one: the configuration with one ‘generalist’ population that could consume all of the pools was able to draw down even the most recalcitrant pools to very low concentrations at depth. For clarity, we illustrate the model solution with the “specialist” configuration below, and illustrate the alternative ecosystem configurations in the Supporting Information (Figs. S12 and S13).

For simplification, we explain and illustrate only the C:N of the solutions in the main text, since the resulting C:P relationships are analogous to those of C:N (see Fig. S2 for the C:P and N:P of the solutions). Though aspects of the resulting N:P of the solutions are plausible, we do not study them here since other processes that we do not explicitly represent must be considered for a robust interpretation of those dynamics (e.g. viruses (Jover et al. 2014)). We choose to illustrate the C:N because the DOC:DON observations (Fig. 1) have a narrower range of uncertainty than those of DOC:DOP (Letscher and Moore 2015), and also because Galbraith and Martiny (2015) emphasize the role of the remineralization of N in controlling the efficiency of the biological carbon pump.

## 3 Results

All three versions of the marine water column model capture typical biogeochemical dynamics: light-driven primary production draws down inorganic nutrients near the surface, and remineralization of sinking POM fills the subsurface with inorganic nutrients (Fig. 4a). Heterotrophic biomass is highest near the surface where production of organic matter is largest (Fig. 4a,c,e).

**Figure 4:**
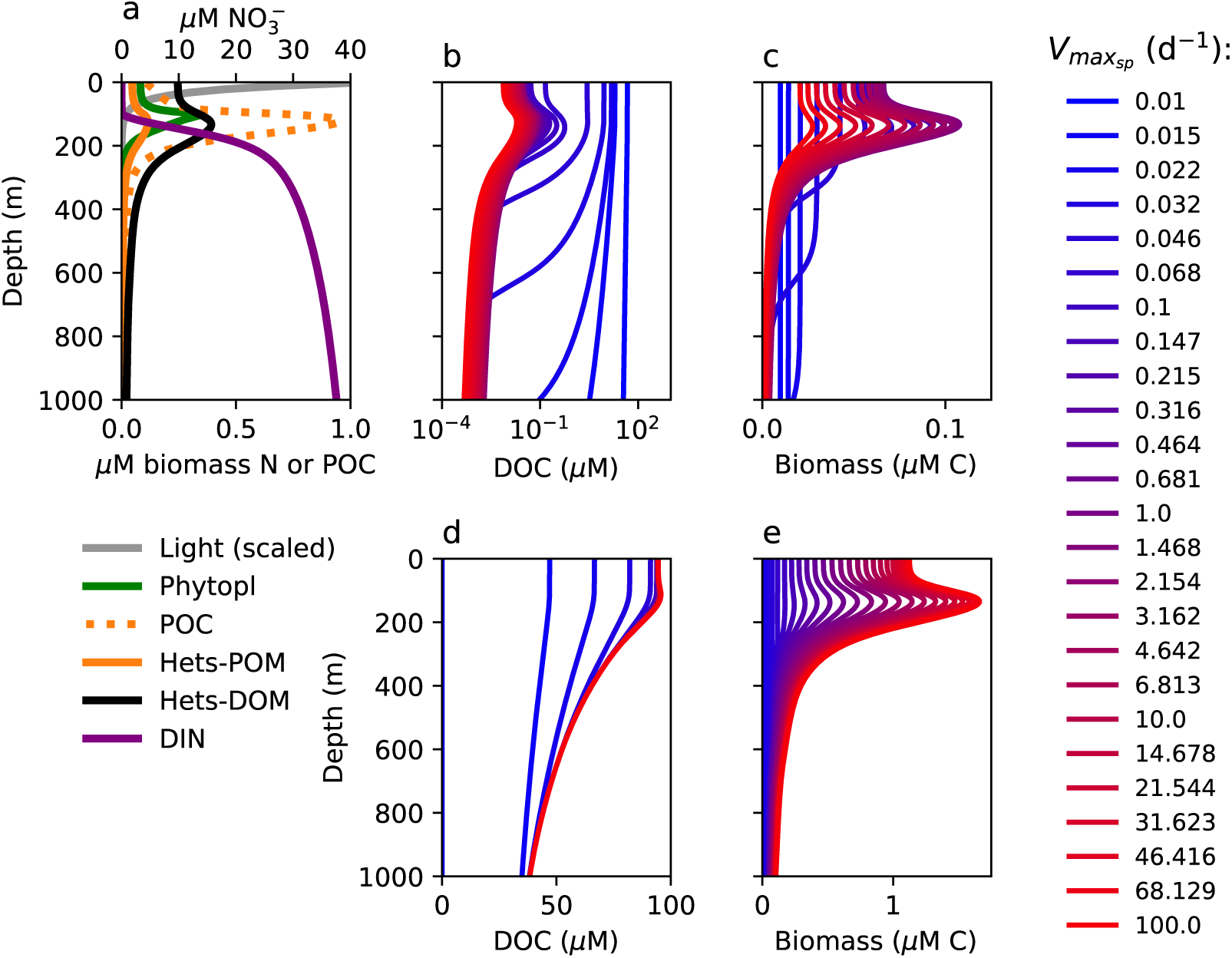
Water column model solutions for (a) light, phytoplankton biomass, POM, DIN, (b and d) concentrations of the 25 classes of DOM, and (c and e) the biomass of the 25 populations of heterotrophs supported by each DOM class. Panels b and c show DOC and biomass concentrations plotted individually, while panels d and e show DOC and biomass concentrations plotted cumulatively, so that the red line indicates total DOC and total biomass concentration. The solutions for Models A, B, and C are visually indistinguishable for these quantities except for the cumulative concentrations of DOC in panel d. Here, output from Model C is shown, but output from all three models is shown in Fig. 1a.

Before presenting specific water column results, we note that in Supporting Text S1 we demonstrate the model’s ability to quantitatively resolve the stoichiometry of aquatic prokaryotic heterotrophic activity by comparing simulations of the metabolic model to two datasets in which the elemental ratio of nutrient supply was varied (Goldman et al. 1987; Godwin et al. 2017). The model captures the observed relationship between the stoichiometry of supply and the stoichiometry of remineralization of labile substrates (Goldman et al. 1987, Fig. S1). Importantly, the model also simulates the observed large increase in C:P of heterotrophic biomass from about 80 to about 600 mol C mol^−1^ P as a function of C:P supply and growth rate (Godwin et al. 2017, Fig. S1). These results provide confidence in meaningful representations of remineralization in the water column model.

### 3.1 DOM concentrations as a function of lability

All versions of the model reproduce two general patterns in DOM concentrations (Fig. 4b,d). First, the modeled concentrations of the more labile DOM classes (the red and purple lines in Fig. 4b,d) are low at the surface (< 0.1 μM C), and decrease to nanomolar concentrations or lower with depth. Second, the concentrations of slower, more “recalcitrant” pools (the blue lines in Fig. 4b,d) are high (1–50 *μ*M C). This DOM accumulates in the model because the supply rates of these DOM classes at the surface exceed their maximum rates of consumption (*V*_*max*_). The timescale of transport (vertical mixing) is also short relative to the consumption rates for these pools, and so mixing distributes the recalcitrant DOM to depth. Intermediate classes that accumulate in the mixed layer are eventually depleted in the mesopelagic (500 and 1000m depths), which is consistent with observations and current conceptual models of DOM dynamics (Hansell 2013). The slowest-degrading pools attain relatively constant concentrations throughout the water column, simulating the ubiquitous concentrations of recalcitrant DOM throughout the ocean at all depths (Hansell 2013).

In the illustrated model versions, elevated DOM concentrations are a result of the accumulation of recalcitrant substrates. However, since the concentration of each labile DOM pool reaches a non-zero minimum, the total concentration of the modeled labile DOM fraction will be a function of the number of pools resolved. This is the mechanism behind the “dilution hypothesis” (Jannasch 1967; Arrieta et al. 2015), which contends that high concentrations of DOM reflect the sum of many low concentrations. Our results are independent of the number of DOM pools resolved because we focus on the stoichiometric patterns and their mechanisms, but they do depend on there being quantitative differences in the consumption rate (i.e. lability) between the pools.

### 3.2 Controls on heterotrophic biomass

Vertical profiles of the 25 heterotrophic biomass ‘clades’ are distinct from the profiles of the DOM class that each consumes (Fig. 4c,e). At the surface, the most abundant heterotrophic clades consume the classes of DOM with the highest rate of supply (the purple lines in Fig. 4c). The clades consuming the most labile substrates are lower in abundance (the red lines) because less of these substrates are produced. Thus supply exerts a strong control on the amount of biomass sustained on labile substrates.

In contrast, at depth, overall biomass is lower, and the majority of biomass is associated with the least labile substrates. These low biomass values, despite abundant resources, can be explained by both top down control (grazing or lysis) and limitation by maximum consumption rate (*V*_*max*_).

We aggregate these multiple controls into an approximation of the steady state concentration of biomass (see Supporting Text S3 for derivation). We then delineate two types of limitations on bacterial biomass: by substrate concentration as “concentration-limited”, and by consumption rate as “*V*_*max*_-limited.” For depleted substrates, because [*DOC*] ≪ *k*_*DOC*_ and 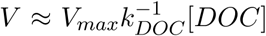, the associated concentration-limited biomass 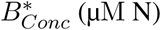 (μM N) can be approximated as:

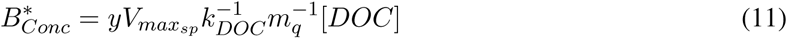

For the recalcitrant classes of DOM, concentrations are high, and so the consumption rate is saturated (*V* ≈ *V*_*max*_). This represents a limitation by the rate of processing by the microbial cells. This processing-limited biomass 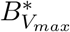 (μM N) can be approximated as:

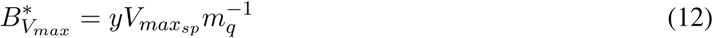

We plot 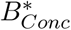 and 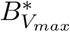 against the modeled biomasses in Fig. 5b (black lines). The concentration-limited contour (dashed black line) matches the modeled biomasses associated with the most labile substrates, and decreases as *V*_*max*_ increases. In contrast, the *V*_*max*_-limited contour (solid black line) matches the biomasses associated with the more recalcitrant DOM classes, and increases with *V*_*max*_. In summary, the model suggests that when a population is limited by the concentration of a substrate, the amount of biomass sustained is proportional to that substrate. When a population is instead limited by the rate at which cells can process the substrate, reflecting the low energy and/or accessibility of the recalcitrant substrate, the amount of biomass sustained is proportional to that rate.

**Figure 5:**
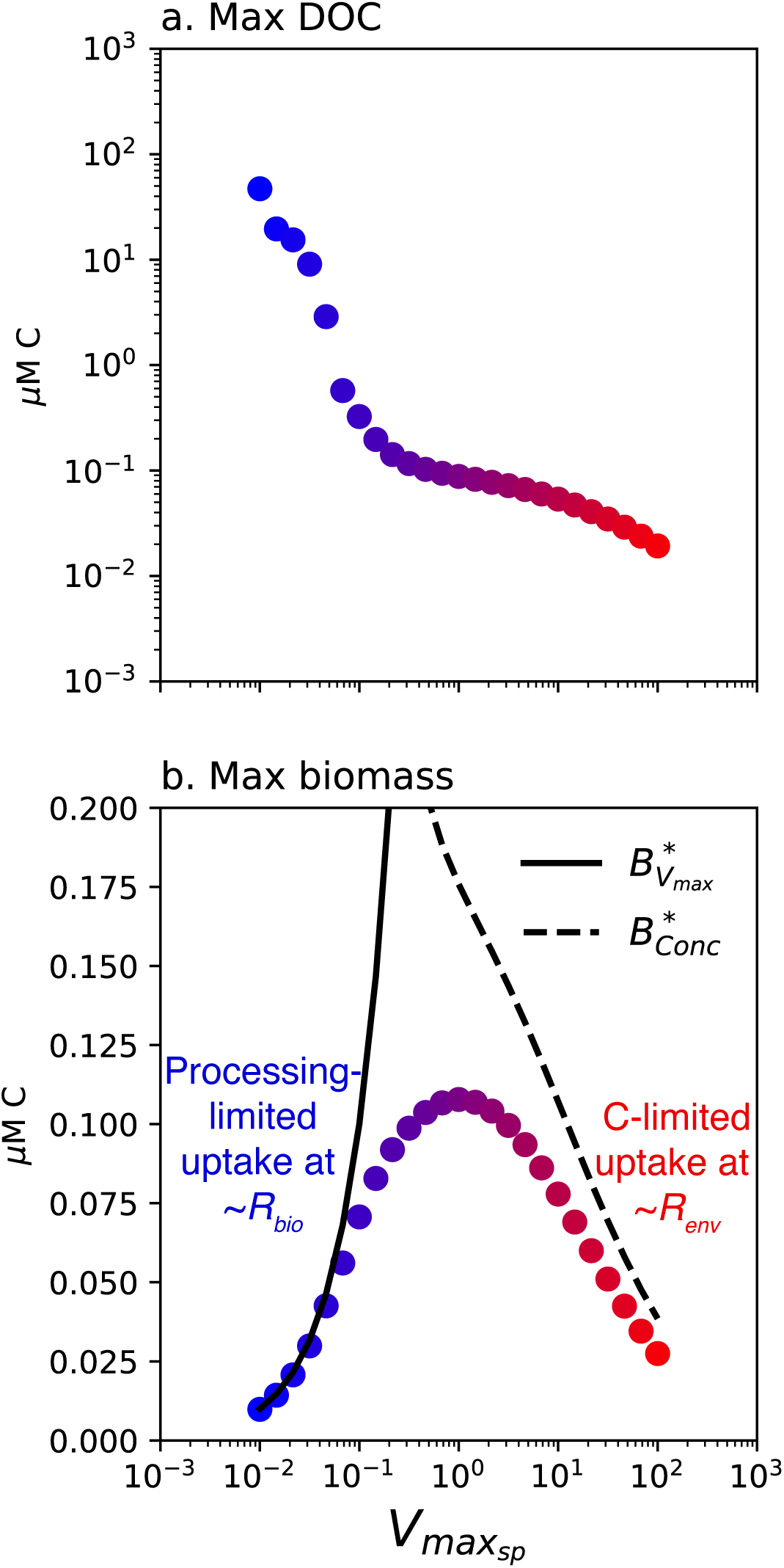
Maximum DOC concentration and maximum biomass concentration as a function of specific maximum uptake rate 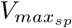 in the water column model. The black lines in panel b indicate the approximations of concentration-limited biomass (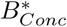; Eqn. 11) and processing-limited biomass (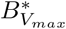 ; Eqn. 12), converted to carbon biomass with *R*_*bio*_.

### 3.3 Enhanced C:N of phytoplankton vs. heterotrophic biomass

The C:N of the biomass pools emerges as a consequence of interactions in the ecosystem model. Despite all populations having the same optimal biomass stoichiometry (*Q*_*minC*_ : *Q*_*minN*_ = 5), C accumulates relative to N in photoautotrophic but not heterotrophic biomass in all model simulations (Fig. 6 first column). Elevated phytoplankton C:N (here to about 15 mol C per mol N) is consistent with observations and models (Goldman et al. 1979; Martiny et al. 2013a,b; Talmy et al. 2016). Modeled heterotrophic biomass C:N remains about 5 throughout the water column (varying from 4–5 in model variations; see Fig. S6 in particular), which is consistent with the mean observed heterotrophic bacterial stoichiometry of 5 ± 1 (Zimmerman et al. 2014). The C:N of the supply of new POM and DOM in the model reflects the weighted average of these biomass pools.

**Figure 6:**
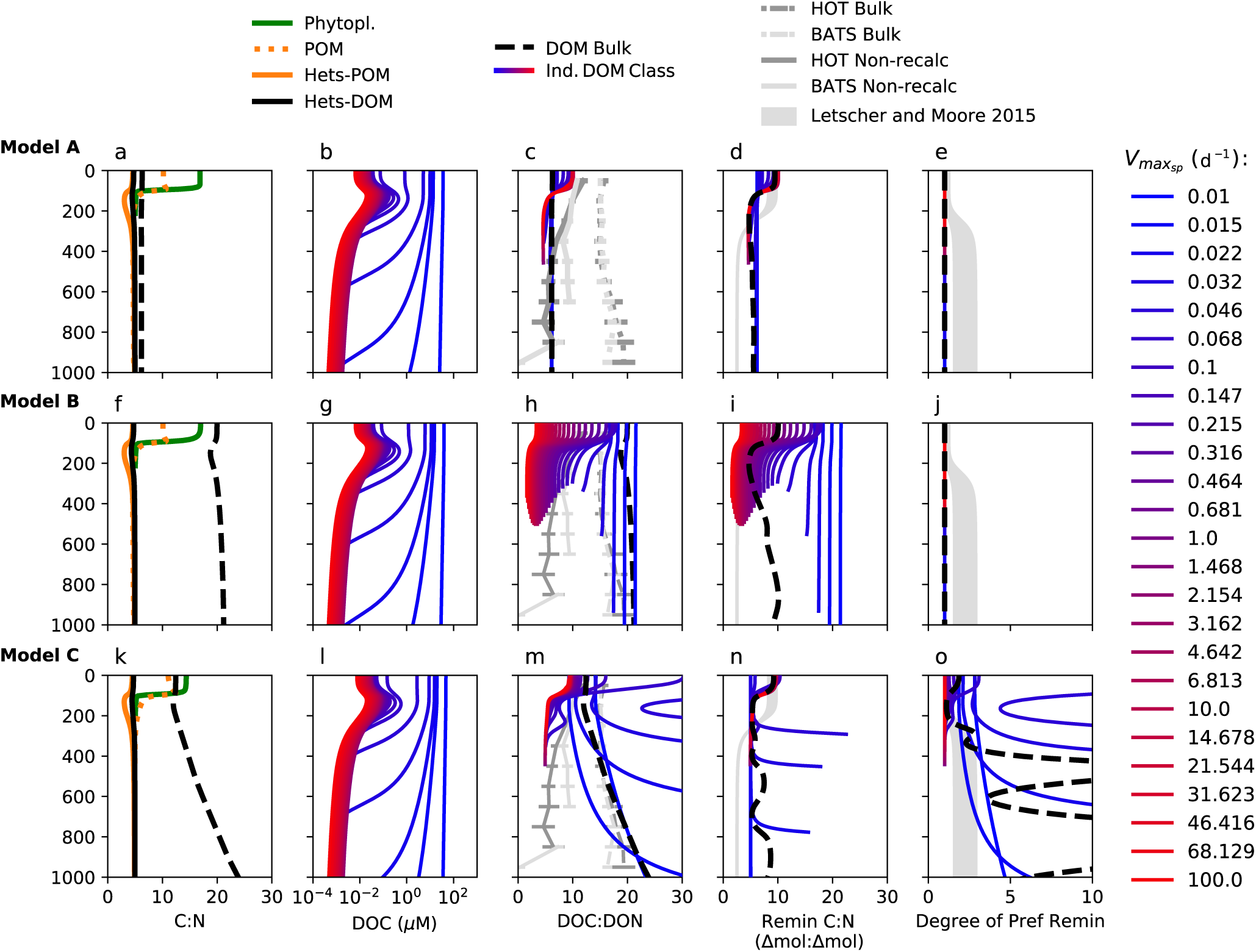
The C:N of the water column model solutions for the three model versions isolating different aspects of control on the stoichiometry of DOM and its remineralization (Models A (a–e), B (f–j), and C (k–o)). Column 1: The C:N of the biomass groups, POM, and bulk DOM. Column 2: the concentration of each of the 25 DOM classes. Column 3: the C:N of each of the 25 DOM classes and of bulk DOM in the model, and the observed C:N of bulk and non-recalcitrant DOM at HOT and BATS from Fig. 1. Column 4: The remineralization ratio of each of the 25 DOM classes and for bulk DOM. Column 5: The degree of preferential remineralization of N vs C for each of the 25 DOM classes and for bulk DOM. In columns 3–5, values for the individual classes are not shown where DOC or DON for that class is depleted below 1 nM. In columns 4 and 5, the gray area represents the envelope of the remineralization ratio for non-recalcitrant substrates and the magnitude of preferential remineralization calculated from observations by Letscher and Moore (2015) (See Supporting Text S4).

**Figure 7:**
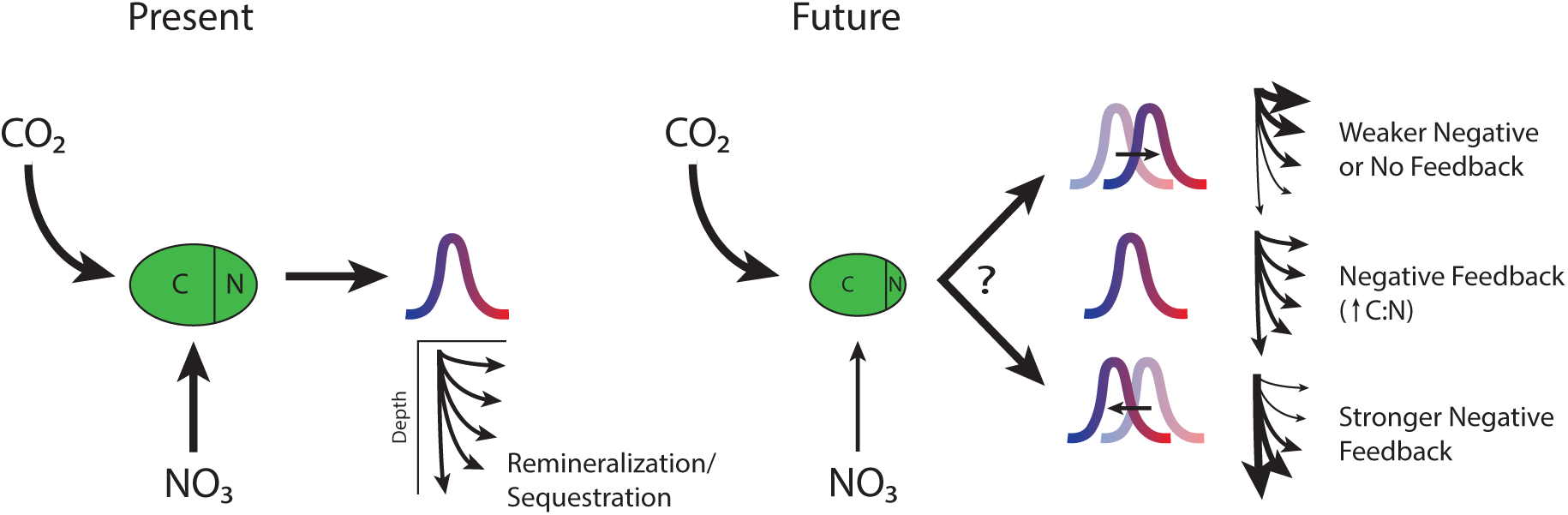
Impact of heterotrophic consumption on a potential climate feedback as a function of changes to the C:N and lability of DOM.

This difference in biomass stoichiometry reflects the difference in the elemental ratios of substrates consumed by phytoplankton vs. heterotrophs. In the sunlit surface, DIC:DIN varies from 2000:0.1 to 2000:0.01, and so when solar energy is abundant and DIN is limiting, phytoplankton fix excess C into organic form, resulting in elevated phytoplankton C:N. In contrast, the modeled C:N of DOM classes varies only from 5 to about 100 mol C per mol N. Our simulation of the dataset of Godwin et al. (2017) in Supporting Text S1 demonstrates how heterotrophic biomass stoichiometry can vary significantly as a result of much larger variations in nutrient supply (up to a 7-fold increase from *R*_*bio*_ in Fig. S1). Thus, while heterotrophs have the potential for significant variation in elemental ratios (Godwin et al. 2017), this potential may not often be observed in the ocean.

The majority of modeled heterotrophs are C-limited, as discussed further below, consistent with what is believed to be true in the ocean. However, if heterotrophic biomass stoichiometry was predominantly influenced by this C-limited state, heterotrophic C:N would be systematically lower than the assigned optimal biomass stoichiometry (*R*_*bio*_). The biomass of the POM-consuming heterotrophs does exhibit this decreased C:N in the subsurface, where POM is most rapidly attenuated by heterotrophs, suggesting that C-limitation may manifest as lower C:N in some locations or for some species.

### 3.4 DOM and remineralization stoichiometry as a function of lability

#### Decreasing C:N of more labile DOM

All three models predict that the C:N of more labile classes of DOM decreases with depth, simulating the observed profiles (Fig. 1c, Fig. 6 third column, red lines). This stoichiometry matches a switch in the C:N of the local supply. In the sunlit surface, more of the DOM supplied originates from C-rich phytoplankton biomass, resulting in elevated C:N. The surface C:N elevation increases with phytoplankton excretion (i.e., with phytoplankton “leakiness”; Fig. S5). Below the surface, more DOM originates from heterotrophic biomass with lower C:N. Because labile DOM is consumed quickly relative to mixing, the C:N of the local supply drives the profile.

Preferential remineralization of one element or the other does not occur for labile substrates (Fig. 6 fifth column, red lines), despite the fact that the remineralization ratios systematically depart from Redfield values. The concentrations of labile DOM are low, and so the uptake stoichiometry (i.e., the remineralization ratio) matches the environmental stoichiometry in all three models (Fig. 6 fourth column, red lines). This is consistent with the findings of Letscher and Moore (2015) of high remineralization C:N at the surface with no preferential remineralization of DOC (Fig. 6 grey shaded areas). Because the heterotrophs deplete the available labile DOC to low concentrations while excreting DIN (see section 3.5), their growth is C-limited.

The gradient in labile DOC:DON and its remineralization ratio is sharper and shallower in the model than in the observations (Fig. 6 third and fourth columns). We attribute this to the simplified representation of POM in the model. Resolving a spectrum of POM classes with different sinking rates would allow for some C-rich phytoplankton biomass to reach deeper depths (due to ballasting or aggregation, for example), resulting in a more gradual transition from high to low C:N. Since we here aim for a mechanistic understanding of heterotrophic elemental cycling rather than accurate predictions, the magnitudes of DOC:DON are approximate, but the patterns are robust. In the real ocean, DOC:DON should additionally be impacted by phytoplankton excretion (as in Fig. S5), decoupling of DON:DOP (not examined here), organic consumption by phytoplankton, and other processes.

#### Enhanced C:N and preferential remineralization of less labile and ‘recalcitrant’ classes

Differences between the three models emerge in the stoichiometry and remineralization ratio of the least labile DOM classes. Because these classes turn over more slowly and accumulate in the water column, the elemental ratios for these classes are impacted by non-local (vertically integrated) DOM supply and uptake stoichiometry.

In Model A, the stoichiometry of consumption matches DOC:DON, and so heterotrophic activity does not affect DOM stoichiometry. The C:N of the recalcitrant classes in these simulations does not increase (contrary to observations), but remains at about 7-8 mol/mol throughout the water column, reflecting an integrated average of heterotrophic and phytoplankton biomass stoichiometry (Fig. 6a). The remineralization ratio matches that of this integrated average (Fig. 6d), and there is no preferential remineralization.

In Model B, the recalcitrant pools are produced with a higher C:N according to our imposed distributions (Fig. 3), and so the C:N of these pools is inherently high (Fig. 6h). The remineralization ratios of these pools are also high because uptake stoichiometry matches the ambient DOC:DON (as in Model A). Therefore, no preferential remineralization occurs in the Model B simulations (Fig. 6j). Here, the C:N of bulk DOM does not increase with depth because the four most recalcitrant classes comprise the majority of the total concentration, and the C:N of these pools reflects our imposed distribution (Fig. 3). It is possible for the C:N of bulk DOM to increase with depth in this model if we were to impose a different (less simple) imposed distribution of DON production. This model does not mechanistically capture the preferential remineralization observed by Letscher and Moore (2015), although it can be thought to represent preferential remineralization implicitly, and thus may be an appropriate description in biogeochemical models without explicit heterotrophic activity.

In Model C, the C:N of uptake for the less labile classes approaches the biologically regulated elemental ratio (*R*_*bio*_) due to the accumulation of these classes to higher concentrations (Fig. 2b). This simulates the targeted consumption of N-rich compounds when the concentration of substrate alone does not limit growth. Growth on these DOM classes is limited by the energy or accessibility of the substrate via the microbial processing rate, and not by C (or N). Since *R*_*bio*_ is lower than the C:N of the ambient stoichiometry (*R*_*env*_), residual DOC remains, and the DOM classes become enhanced in DOC (Fig. 6m). This constitutes preferential remineralization (Fig. 6o).

Intermediately recalcitrant DOM classes (blue-purple lines) accumulate higher in the water column but are eventually depleted at depth. These classes reach the highest C:N values in the model (over 30 mol/mol; Fig. 6m,n), and are associated with the strongest degree of preferential remineralization (Fig. 6o). The heterotrophs consuming these classes of DOM are controlled by either 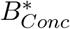 or 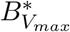, depending on the location within the water column. In the surface, the biomass is limited by the processing rate 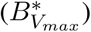, and so preferential remineralization strips out DON relative to DOC. At depth, where supply decreases, the biomass depletes these intermediate pools and becomes limited by concentration 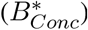. As a result, below the euphotic zone where DOM supply rates drop, the intermediate DOM pools experience high rates of preferential remineralization, yielding high C:N. This formation of high C:N DOM is consistent with the “low-nutrient,” carbon-rich organic matter discussed by (Fawcett et al. 2018), which we discuss below.

In Model C, the bulk (average) degree of preferential remineralization of N increases with depth (Fig. 6o). Therefore, Model C suggests that the C:N of bulk DOM may increase not solely because of a progressively lower proportion of labile DOM, but also because the ‘recalcitrant’ classes themselves become increasingly enhanced in carbon (Fig. 6m). The bulk degree of preferential remineralization of N oscillates significantly from high to low values, which is a consequence of the discretization of the DOM classes in the model. This behavior is due to the strong increase in the preferential remineralization of N for individual classes as each nears its depletion depth (as described above), rather than a more steady continuum of depletion that we would expect in the real ocean.

Though qualitatively robust, the magnitudes of the C:N enhancement and the increase in bulk DOC:DON with depth in Model C exceed the observed magnitudes (Fig. 6m,o). This may indicate that some less labile substrates are taken up at ratios that match their *in situ* ratios. The modeled magnitudes are also sensitive to the balance of DOM supply vs. consumption, which are affected by many of parameter values as well as the community consumption configuration. For example, the increase in DOC:DON with depth was enhanced with lower microbial mortality rates and muted for lower maximum carbon yields for the populations consuming the accumulated substrates (Figs. S7 and S11). Because these magnitudes are consequences of many processes, explicitly representing heterotrophic dynamics is necessary for mechanistically capturing the degree of preferential remineralization in the model.

In summary, Models B and C both predict elevated C:N for less labile and recalcitrant DOM classes. Model A does not allow for elevated C:N for these classes, and thus cannot describe the real ocean unless all of the ‘excessive’ recalcitrant DOC (the amount of DOC that results in the elevated C:N) is from non-marine-biomass sources, such as abiotic photodegradation/phototransformation, black carbon, or terrestrial carbon. Critically, only Model C predicts preferential remineralization consistent with observed patterns (Letscher and Moore 2015). Thus, we infer that some degree of biologically regulated consumption characterizes DOM remineralization.

### 3.5 Heterotrophic DIN assimilation

In all three model versions, the heterotrophs assimilate DIN in addition to DOM, and excrete DIC, DIN, and DIP. Thus, they simultaneously assimilate and remineralize 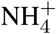 as has been previously observed (Tupas and Koike 1990; Tupas et al. 1994). In the surface, the heterotrophic biomass consuming labile substrates takes up DIN at nearly equal rates as DON (Fig. S14), since the C:N of labile substrates is nearly 2*R*_*bio*_. This rate of uptake is also nearly equal to the rate of phytoplankton DIN uptake in all three models.

However, since most of the DOC consumed is oxidized for energy production, the majority of heterotrophs consume N in excess because their growth is either C-limited or processing-rate-limited, whether taking up DIN or not. Therefore, when DIN assimilation is allowed, DIN excretion increases accordingly (Fig. S14), resulting in the same net N retention in biomass as theorized by Kirchman (1994). Thus, the model suggests that phytoplankton do not compete with heterotrophs for DIN in a steady state environment. When DIN assimilation is not allowed, rates of primary production are nearly identical (< 1% difference).

In Model C, however, the uptake of DIN by heterotrophs does impact total DOC concentrations, which indicates that some heterotrophic growth is N-limited in the water column. When DIN assimilation is not allowed, recalcitrant DOC reaches higher concentrations, and total DOC is about 20% higher (Fig. 1), despite primary productivity remaining the same. (In Models A and B, DIN assimilation has no impact on DOC concentrations or productivity.) The model predicts that the growth of the heterotrophs consuming intermediately labile DOM classes (purple lines) becomes N-limited in the subsurface. As this intermediately labile DOM is eventually depleted to low concentrations at depth, the C:N of these classes becomes high from the preferential extraction of DON (higher than 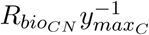), as discussed above. DIN uptake supplements heterotrophic growth in these conditions. This captures the findings of Fawcett et al. (2018), who observed heterotrophic uptake of nitrate within and below the nitricline, and concluded that this uptake supplemented the remineralization of N-poor organic matter.

## 4 Discussion

We develop an ecosystem model with mechanistic descriptions of the transformations of carbon, nitrogen, and phosphorus to and from organic form by the production and respiration of microorganisms. Model solutions provide mechanistic explanations for observed marine global biogeochemical patterns: the differences in the C:N of phytoplankton vs. heterotrophic biomass and changes in DOC:DON and remineralization ratios with depth below the sunlit zone. The framework links microbial cellular metabolism to global-scale biogeochemistry, improving our understanding of how ecological dynamics mediate ocean C storage.

The water column model predicts the distinct stoichiometries of nutrient-limited phytoplankton biomass (higher C:N; Goldman et al. 1979; Martiny et al. 2013a; Talmy et al. 2016) and C-limited heterotrophs (lower C:N; Zimmerman et al. 2014). This is the first time to our knowledge that the contrasting dynamics governing phytoplankton vs. heterotrophic biomass stoichiometry have been explored in a model. In all model versions, heterotrophic biomass stoichiometry remains relatively constant because heterotrophs experience C:N and C:P supply ratios that are relatively constant compared to photoautotrophs that can utilize the large pool of DIC. The relatively lower, constant C:N of heterotrophic biomass is not a model artifact, as we demonstrate that simulated heterotrophs do have the capability to drastically vary their biomass stoichiometries with varying substrate supply (Fig. S1). Thus the model reconciles the understanding that heterotrophic biomass stoichiometry has the potential to vary widely but largely remains consistent in marine environments.

Each of three non-mutually exclusive models of uptake stoichiometry represent an aspect of real control on DOM remineralization. Two of these controls – higher lability of DON (Model B) and biologically regulated uptake (at a lower C:N than supplied; Model C) – capture the observed elevated C:N of bulk DOM. All models simulate the C-limited growth of heterotrophs on more labile substrates, which constitutes the bulk of remineralization in the surface layer.

Only Model C results in preferential remineralization of N. This resolves the apparent paradox by suggesting that C-limited organisms 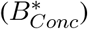 do not exhibit preferential remineralization, while those that are processing-limited 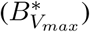 do target N-rich compounds (or fragments of compounds). In other words, we hypothesize that preferential remineralization can only be understood when considering both the stoichiometry of consumption and the controls on heterotrophic biomass. This mechanism is consistent with observations that microorganisms can regulate the stoichiometry of their substrate consumption (You et al. 2013; Burkhardt et al. 2014). For example, Burkhardt et al. (2014) show that the C:P of remineralization remains constant despite variation in the C:P of organic supply.

Model C qualitatively, and somewhat quantitatively, reproduces the pattern and the degree of preferential remineralization computed by Letscher and Moore (2015) (Fig. 6o). Thus we have successfully linked microbial-scale processes of substrate uptake with bulk water column measurements. The degree of preferential remineralization in the model is higher at depth than the observations suggest. This may indicate that reality does indeed reflect a combination of the three “end-members” governing organic uptake stoichiometry (i.e., Models A, B, and C), and thus that uptake of some of the less labile substrates does match environmental stoichiometry. It may also reflect that the supply of more labile substrates to the deep ocean is underestimated due to the simplified parameterization of the POC flux.

Model C modifies the explanation of Letscher and Moore (2015) and Aminot and Kérouel (2004) for the increase in the C:N of bulk DOM with depth. Results suggest that this increase may not only reflect a decrease in the average lability of DOM, as in the previous explanation, but also an increase in the C:N of more recalcitrant DOM itself as preferential remineralization increases in strength.

Furthermore, Model C may serve as a useful parameterization for DOM consumption in global biogeochemical models because of its realistic treatment of stoichiometry across the DOM lability spectrum. At low concentrations, uptake ratios match supply ratios, realistically simulating the consumption of depleted, labile substrates and C-limited growth. At high concentrations, uptake ratios converge to the biologically regulated ratio, realistically simulating the targeted consumption of the N- and P-rich portions of less labile compounds where growth is limited by processing rate.

In the model, the C:N of both the newly produced DOM and POM reflects the weighted average of the biomass pools (Fig. 5). Since we resolved only one pool of POM with a lability that matches the mean lability of DOM, the C:N of non-recalcitrant DOM and POM are nearly equal. However, in the ocean, the C:N of non-recalcitrant DOM and POM may be decoupled, resulting in lower surface POC:PON than DOC:DON (Martiny et al. 2013b). Specifically, phytoplankton excretion of DOC will increase the C:N of surface DOM (when their *γ >* 0 as in Fig. S5), and phagotrophic consumption by zooplankton will decrease the C:N of POM (Talmy et al. 2016). Both of these processes will decrease POC:PON relative to DOC:DON, and contribute to a relatively lower C:N of the DOM supplied from POM. This will primarily impact the labile DOC:DON, since it is driven by local supply, but will not change the dynamics outlined here.

In reality, POM should also exhibit a spectrum of lability, allowing a pattern of increasing C:N with depth analogous to DOM (Schneider et al. 2003), and with the less labile portions reaching the bottom of the ocean and forming the sediment. In the present model, the DOM production stoichiometry is not altered by its pathway through the POM pool because POM is not subject to preferential remineralization. If a spectrum of POM lability were represented, preferential remineralization of PON would perhaps decrease the C:N of DOM sourced at depth, if relatively more DON than DOC is released from the POM, though this does not qualitatively change the proposed controls on DOM stoichiometry.

Results suggest that phytoplankton and heterotrophic bacteria do not compete for DIN because heterotrophic growth is predominantly C-limited in the sunlit layer. If heterotrophs are allowed to assimilate DIN in the euphotic zone, they excrete DIN at higher rates as a consequence. This provides a mechanistic explanation for simultaneous assimilation and remineralization of 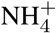 observed by (Tupas and Koike 1990) and (Tupas et al. 1994). However, the model also predicts N-limited growth of some heterotrophs below the sunlit layer. This is consistent with the observations of nitrate-assimilation by N-limited heterotrophs consuming intermediately labile, “nutrient-poor” DOM in the subsurface (Fawcett et al. 2018).

Critically, our results suggest how remineralization may impact changes to the efficiency of the biological and/or microbial carbon pumps. Variable remineralization stoichiometry may modify the negative climate feedback proposed by Galbraith and Martiny (2015). In the proposed feedback, C sequestration is enhanced due to increased C:N of organic matter produced by phytoplankton at low latitudes under future climate scenarios. Our results suggest that the strength of this negative feedback is dependent on the associated stoichiometry of remineralization, and cannot be simply estimated in a model that assumes constant Redfieldian stoichiometry. One can envision three scenarios that all result in an increase in the C:N of primary production: 1) the C:N of all organic pools increases uniformly, 2) the production of high C:N labile organic matter (e.g. sugars) increases while the rest of the pools maintain a lower C:N, or 3) the production of high C:N recalcitrant organic matter (e.g. cell wall) increases while the rest of the pools maintain a lower C:N. Based on the hypothetical relationship between lability and remineralization stoichiometry that emerges from our model, we suggest that the signal of increased C:N (the basis of the negative feedback) will only persist if scenario 1 or 3 occurs. In scenario 2, the remineralization ratio will increase with the labile C:N accordingly, and so this fraction will not contribute to increased C storage. Moreover, scenario 3 should result in a stronger negative feedback than originally proposed because the degree of preferential remineralization is stronger for recalcitrant pools, and so carbon would be retained in the system for longer periods of time than would have occurred with constant remineralization stoichiometry.

While heterotrophic community structure was not the focus of this work, the model does provide some insight into large-scale patterns of heterotrophic function and constraint on how these processes should be represented in global-scale models. Specifically, we find that the patterns of DOM accumulation are robust across many different configurations of community consumption as long as a sufficient penalty for consuming multiple substrates was incorporated. The exception is when a large aggregation of heterotrophic biomass, modeled as one single “generalist” population, is allowed to consume all classes of DOM. Despite equal specific rates of DOM uptake as in the other configurations, the large biomass of this aggregation enables higher volumetric rates of recalcitrant DOM consumption, preventing the accumulation of this pool throughout the water column. These results suggest that the populations that degrade more recalcitrant substrates are specialized to do so. This is consistent with recent observations that suggest ‘rare’ taxa are responsible for degrading more specialized substrates (Rivett and Bell 2018), and that the ability to degrade high molecular weight substrates is a more specialized, less widely distributed trait than that of low molecular weight degradation (Logue et al. 2016; Balmonte et al. 2019).

The model developed here constitutes progress towards the predictive framework called for by Galbraith and Martiny (2015) to systematically address stoichiometric flexibility in marine ecosystems in order to evaluate its control of global biogeochemical cycling. Due to the focus on prokaryotes, we do not address other factors that affect the stoichiometry of organic substrates. For example, Talmy et al. (2016) demonstrate that phagotrophic zooplankton, which have a biomass C:N more similar to heterotrophic bacteria than to phytoplankton, may lower the C:N of the particulate organic pool by respiring proportionally more C than N. Additional work should also address the impact of viral lysis on marine stoichiometry, and in particular the decoupling of N and P cycling (Jover et al. 2014).

## 5 Conclusions

Much attention has been given to the stoichiometry of primary production as a function of inorganic nutrient supply to the ocean surface. Here, we additionally resolve the stoichiometry of the “reverse” reaction – the remineralization by heterotrophs – by resolving the elemental flow of uptake, growth, and excretion by heterotrophs in addition to phytoplankton in an ecosystem model. We demonstrate how their interactions reproduce globally comprehensive patterns in the stoichiometry of dissolved organic matter and its remineralization.

This work provides insight into how heterotrophic growth and respiration mediates the stoichiometry of organic matter in the ocean, with implications for how this remineralization may ultimately impact the efficiency of the biological carbon pump. The model developed here can be combined with stoichiometric resolution of zooplankton activity, resolution of nitrogen fixation and denitrification, and other relevant biogeochemical processes, and probed to study how biologically driven C storage may change globally with changes to the environment locally, such as changes in nutrient supply to the oligotrophic surface ocean.

## 6 Acknowledgements

We thank R. Letscher for providing the DOM data compilations, and C. Godwin for providing the datasets from the chemostat experiments. EJZ was supported by the Simons Foundation (Postdoctoral Fellowship in Marine Microbial Ecology). EJZ and NML were supported by the Simons Foundation: The Simons Collaboration on Principles of Microbial Ecology (PriME #542389). The ecosystem model code is available at https://zenodo.org/record/3497551 (DOI: 10.5281/zenodo.3497551).

