## Supporting Information for "Systematic variation in marine dissolved organic matter stoichiometry and remineralization ratios as a function of lability"

**Contents:**

1. Text S1 to S4
2. Figures S1 to S14
3. Tables S1 to S2

### Overview

In Supporting Text S1, we describe the details of the comparisons of the metabolic model with two experimental datasets (Fig. 1, Table 1). In Supporting Text S2, we describe the details, equations, and parameters for the ecosystem model (Table 2). Fig. 2 illustrates the C:N, C:P, and N:P of the model solutions.

Figs. S3–S13 illustrate and explain some of the parameter sensitivity experiments, including the different community consumption configurations mentioned in the main text. Plausible variations in all parameter values affect solutions quantitatively but not qualitatively. For example, while water column profiles remained consistent, increasing $y_{maxC}$ increases microbial biomass and decreases DOM concentrations; increasing uptake affinity ($V_{max_S} k_S^{-1}$) decreases concentrations of the more labile ("concentration-limited") pools; and increasing mortality rates increases DOM concentrations and growth rates and decreases biomass. Many of these parameters affect the environmental fitness of the heterotrophic populations, and so these differences become important for studies of the controls on the functional diversity and community structure of real heterotrophic populations. Importantly, varying the $Q_{min}$ ratios ($Q_{minC} : Q_{minN}$) gives proportionally varying values of C:N across all pools, although the resulting relationships between pools remain the same across the variations.

Fig. 14 provides a detailed comparison between models with and without heterotrophic DIN assimilation. Supporting Text S3 provides detail for the derivation of the steady-state biomass approximations described in the main text. Finally, Supporting Text S4 explains the estimate of the envelope of the remineralization ratio and degree of preferential remineralization calculated from observations (corresponding to the gray shaded areas in Fig. 5 in the main text).

### Text S1: Comparison with experimental datasets

**Goldman et al. (1987)** We tested the ability of the metabolic model to quantitatively predict the stoichiometry of remineralized inorganic nutrients using the results of Goldman et al. (1987) (their Experiment A). Goldman et al. (1987) quantified the biomass accumulation and excretion of natural assemblages of marine bacteria grown in a batch culture with varying combinations of argenine and glucose to alter the C:N of organic substrate supply.

To match the experimental setup, we ran the model as a virtual batch culture forced with a range of C:N of initial organic supply, matching the ratios used by Goldman et al. (1987). $PO_4^{2+}$ was also supplied following the experimental protocol. Given that argenine and glucose are considered as labile substrates, we supplied the chemostat with one pool of labile DOC and one pool of labile DON. (Thus the distinctions between Models A, B, and C described in the main text are irrelevant.) We found that the specific maximum uptake rate required to simulate the fast consumption of the labile substrates was on the order of 10 $d^{-1}$ (Table 1). The maximum carbon yield $y_{maxC}$ was set to 0.6 mol biomass C synthesized per mol DOC consumed, and the maximum growth rate was set to 2 $d^{-1}$. Other metabolic parameters were the same as in the water column model (Table 2). The additional accumulation of $NH_4^+$ and $PO_4^{3-}$ from lower rates of respiration could be simulated either by respiration of a less labile pool, sourced from microbial biomass waste, or by assuming a maintenance cost. In the present version, we assumed the former, with less labile DOM sourced from the loss rates of the microbial population.

The model captured the decrease in the excreted $NH_4^+$ and $PO_4^{3-}$ with the increase in the C:N of supply. This reflects the capability of the model heterotrophic population to utilize C and N with different efficiencies in accordance with Eqn. 5 in the main text. When the growth of the population was N-limited (C:N = 10: the red line in Fig. 1 A–C), the population used the organic N with close to 100% efficiency, and no $NH_4^+$ accumulated initially as observed. After the heterotrophs had consumed the labile organic supply during the exponential growth phase (after about two days), consumption and excretion at lower rates resulted in later accumulation of $NH_4^+$ and $PO_4^{3-}$, as observed. Overall, the model simulated how the stoichiometry of supply dictates the stoichiometry of remineralization of labile substrates.

**Godwin et al. (2017)** We next tested the ability of the model to capture changes in biomass stoichiometry. We compared the model to experiments by Godwin et al. (2017) with a single strain of aquatic bacteria in a chemostat with large variations in C:P input (Fig. 1 D–E). Godwin et al. (2017) examined the growth of *Brevundimonas* (among other strains, which showed similar patterns) as a function of two factors: i) the C:P of supply, altered by changing the ratio of glucose and phosphate input, and ii) the relative growth rate (relative to the maximum rate) $\mu : \mu_{max}$, altered by changing the dilution rate of the chemostat.

Following the experimental methods, we ran the model as a virtual chemostat with the incoming concentration of glucose (as one DOC pool) set to 23.88 mmol C $L^{-1}$. Inorganic N was supplied in abundance (as $NH_4^+$ with a C:N of 1.3:1), and $PO_4^{3-}$ was supplied to give incoming C:P ratios of 100:1, 316:1, 1000:1, 3162:1, and 10,000:1. Dilution rate $D$ was set at varying fractions of the maximum growth rate of *Brevundimonas* as indicated in Fig. 1. The model parameters were the same as in the water column model except for: i) the maximum growth rate $\mu_{max}$, which was set to that of *Brevundimonas* (0.159 $h^{-1}$), ii) the maximum specific rate of uptake of DOC $V_{max_{sp}}$, which was set to 10 $d^{-1}$ to match the observed growth, and iii) the maximum carbon yield $y_{maxC}$, for which a value of 0.8 best reproduced the observed variations in growth efficiencies (Table 1). The effective carbon growth efficiency $y_C$ (Fig. 1E) was calculated from the rate of carbon biomass synthesis $mu_C Q_C$ and DIC excretion as:

$$y_C = \frac{\mu_C Q_C}{\mu_C Q_C + E_{DIC}} \tag{1}$$

The model captured the observed increase in C:P of biomass and the decrease in bacterial growth efficiency (the effective carbon yield $y_C$) with increasing C:P of nutrient availability and decreasing $\mu : \mu_{max}$ (Fig. 1 D–E). This anticipates that the flexibility of biomass stoichiometry decreases as a particular organism's growth rate increases because of the saturating relationship between growth rate and cell quota (Eqn. 1) (Klausmeier et al. 2004; Godwin et al. 2017). Critically, the C:P of the heterotrophic biomass varied from about 80 to about 600 mol C $mol^{-1}$ P.

**Equations** The batch cultures with argenine and glucose additions was modeled as an initial value problem. Each tracer $C$ was a function of sources and sinks $S_C$ as:

$$\frac{\partial C}{\partial t} = S_C \tag{2}$$

The sources and sinks were as for the ecosystem model (Supporting Text S2), except that only two classes of DOM were resolved. One class represented the additions of the labile pools as described above, and the other represented the DOM formed from the mortality of the biomass and remineralized at a slower rate (matching the POM and the mean of the DOM remineralization rates in the ecosystem model).

For the virtual chemostat model of the *Brevundimonas* population, each tracer $C$ was a function of sources and sinks $S_C$, as well as the dilution rate $D$ and incoming concentration $C_{in}$ as:

$$\frac{\partial C}{\partial t} = D(C_{in} - C) + S_C \tag{3}$$

Only DOC, $NH_4^+$, and $PO_4^{3-}$ were supplied by setting incoming concentration $C_{in}$ as described above. The sources and sinks were as for the ecosystem model (Supporting Text S2) except that the mortality of the populations was not included. (Additional mortality was negligible given the loss rate from the chemostat dilution rate).

### Text S2: Water column model detail

**Equations** A total of 197 state variables are resolved in the 1D vertical water column model: 100 describe the 25 populations of the DOM-consuming heterotrophic community, 75 describe the 25 pools of DOM, 7 describe POM and the POM-consuming heterotroph, 4 describe the phytoplankton population, 6 describe the two nitrifying populations, and 5 describe inorganic nutrient concentrations (DIC, $NH_4^+$, $NO_2^-$, $NO_3^-$, and $PO_4^{2-}$). Total carbon, nitrogen, and phosphorus are each conserved over the domain.

Each tracer $C$ is vertically diffused throughout the water column as function of the diffusive coefficient $\kappa$ as:

$$\frac{\partial C}{\partial t} = \nabla \cdot (\boldsymbol{\kappa} \nabla C) + S_C \tag{4}$$

where $S_C$ are additional sources and sinks.

Phytoplankton and heterotrophs are modeled with the Droop formulation, represented below as functional type $i$. For each type, total uptake $V_{i,j}$ of element $j$ is the sum of the uptake of one or more substrates $S$ containing element $j$. (Here for clarity, we write the uptake of each substrate as $V_{i,S}$.) Each DOC, DON, and DOP pool $k$ (where $k$ = 1–25) is differentiated by a specific maximum uptake rate, and is produced according to the distribution in Fig. 3 with weight $\omega_k$. Two additional nitrifying microorganism functional types are included following the parameterization in Zakem et al. (2018), resolving nitrogen-based biomasses of ammonia-oxidizing organisms $B_{AOO}$ and nitrite-oxidizing organisms $B_{NOO}$. Table 2 lists all parameters, their dimensions, and values. For each tracer, sources and sinks (as well as advection POM with vertical velocity $w_s$) are as follows:

$$S_{X_i} = \mu_i X_i - L_i X_i \tag{5}$$

$$S_{B_{i,j}} = V_{i,j} X_i - E_{i,j} X_i - L_i B_{i,j} \tag{6}$$

$$S_{B_{AOO}} = \mu_{AOO} B_{AOO} - L_{AOO} B_{AOO} \tag{7}$$

$$S_{B_{NOO}} = \mu_{NOO} B_{AOO} - L_{NOO} B_{NOO} \tag{8}$$

$$S_{DOC_k} = \frac{\omega_k}{\sum_{\omega_k}} \left[ f_{mort} L_i B_i + (\alpha - 1) V_{i,POC} X_i + E_{i,DOC} X_i \right] - V_{i,k} X_i \tag{9}$$

$$* \, S_{DON_k} = \frac{\omega_k}{\sum_{\omega_k}} \left[ f_{mort} L_i B_i + (\alpha - 1) V_{i,PON} X_i + E_{i,DON} X_i \right] - V_{i,k} X_i \tag{10}$$

$$* \, S_{DOP_k} = \frac{\omega_k}{\sum_{\omega_k}} \left[ f_{mort} L_i B_i + (\alpha - 1) V_{i,POP} X_i + E_{i,DOP} X_i \right] - V_{i,k} X_i \tag{11}$$

$$S_{POC} = (f_{mort} - 1) L_i B_i - \alpha V_{i,POC} X_i - \frac{\partial (w_s POC)}{\partial z} \tag{12}$$

$$S_{PON} = (f_{mort} - 1) L_i B_i - \alpha V_{i,PON} X_i - \frac{\partial (w_s PON)}{\partial z} \tag{13}$$

$$S_{POP} = (f_{mort} - 1) L_i B_i - \alpha V_{i,POP} X_i - \frac{\partial (w_s POP)}{\partial z} \tag{14}$$

$$S_{DIC} = \sum_i E_{i,DIC} X_i - \sum_i V_{i,DIC} X_i - R_{bio_{CN}} \mu_{AOO} B_{AOO} - R_{bio_{CN}} \mu_{NOO} B_{NOO} \tag{15}$$

$$S_{\mathrm{NH}_4^+} = \sum_i E_{i,\mathrm{NH}_4} X_i - \sum_i V_{i,\mathrm{NH}_4} X_i - y_{\mathrm{NH}_4}^{-1} \mu_{AOO} B_{AOO} - \mu_{NOO} B_{NOO} \tag{16}$$

$$S_{\mathrm{NO}_2^-} = - \sum_i V_{i,\mathrm{NO}_2} X_i - (y_{\mathrm{NH}_4}^{-1} - 1) \mu_{AOO} B_{AOO} - y_{\mathrm{NO}_2}^{-1} \mu_{NOO} B_{NOO} \tag{17}$$

$$S_{\mathrm{NO}_3^-} = - \sum_i V_{i,\mathrm{NO}_3} X_i - y_{\mathrm{NO}_2}^{-1} \mu_{NOO} B_{NOO} \tag{18}$$

$$S_{\mathrm{PO}_4^{2-}} = \sum_i E_{i,\mathrm{PO}_4} X_i - \sum_i V_{i,\mathrm{PO}_4} X_i - R_{bio_{NP}^{-1}} \mu_{AOO} B_{AOO} - R_{bio_{NP}^{-1}} \mu_{NOO} B_{NOO} \tag{19}$$

*In Model C, DON and DOP have different distributions, so production of each pool follows a different weighting than production of DOC pools, and so $\frac{\omega_k}{\sum_{\omega_k}}$ is replaced by $\frac{\omega_{NPk}}{\sum_{\omega_{NPk}}}$.

Growth rates $\mu_{i,j}$, uptake rates $V_{i,j}$, and excretion rates $E_{i,j}$ (written to indicate the substrate being excreted in the above equations) for each microbial population (except for the nitrifiers) are calculated with Eqns. 3, 4, and 5, and modified by temperature with $\gamma_T$ (Eqn. 28; Table 2). Heterotrophs are allowed to take up both organic and inorganic forms of N and P, and phytoplankton take up all species of DIN, as described below.

**DOM production** DOM was produced from the 'sloppy feeding' on POM and/or directly from the mortalities of the biomass populations. This DOM supply was partitioned into the 25 classes following a lognormal distribution. The logarithmic mean rate of 1 $d^{-1}$ reflects average marine heterotrophic bacterial growth rates and efficiencies (Robinson and Williams (2005); Kirchman (2016)). The width of the distribution was set as $\sigma = 2$, which is similar to a ballpark estimate for the distribution of remineralization rates in the ocean ($\sigma = 2.4$; Rothman (2014)), and which allowed for the simulation of realistic differences in surface and deep [DOC] and DOC:DON.

**Heterotrophic uptake of DIN and DIP** Heterotrophs were allowed to take up DIN and DIP. A maximum uptake rate of DIN and DIP for each population was calculated using the same kinetic parameters as for phytoplankton uptake. The actual uptake was then calculated as the minimum of this maximum uptake rate and the amount of N or P 'deficit' in DOM consumption. This deficit was the difference between the C:N and C:P of DOM consumption and $R_{bio_{CN}}$ and $R_{bio_{CP}}$, respectively.

**Phytoplankton** The model allows for excess C fixation by phytoplankton. We assume that the maximum rate of DIC uptake for C fixation by phytoplankton is proportional to the maximum rate of DIN uptake as $R_{bio_{CN}}$. However, allowing phytoplankton to take up DIC at a maximum rate resulted in unrealistic overabundance of the phytoplankton C quota. Therefore, for phytoplankton only, we allowed uptake $V_S$ to be moderated by maximum quota $Q_{max_j}$ (mol cell$^{-1}$), following Thingstad (1987) and Verdy et al. (2009), as:

$$V_S = V_{max_S} \left( \frac{Q_{max_j} - Q_{i,j}}{Q_{max_j} - Q_{min_j}} \right) \frac{S}{S + k_S} \tag{20}$$

for each substrate $S$ containing element $j$.

Following Follows et al. (2007) and Dutkiewicz et al. (2015), phytoplankton grow as a function of a maximum growth rate $\mu_{max}$ ($d^{-1}$), with type-specific limitation by nutrients ($\gamma_{N_i}$), type-specific limitation by light ($\gamma_{I_i}$), and modification by temperature ($\gamma_T$) as:

$$\mu_{P_i} = \mu_{max P_i} \gamma_{N_i} \gamma_{I_i} \gamma_T \tag{21}$$

Nutrient limitation is a function of the total concentration of all species of DIN:

$$\gamma_{N_i} = \min\left[1, \frac{\mathrm{NH_4^+}}{\mathrm{NH_4^+} + k_{\mathrm{NH_4}}} + \frac{\mathrm{NO_2^-}}{\mathrm{NO_2^-} + k_{\mathrm{NO}_x}} + \frac{\mathrm{NO_3^-}}{\mathrm{NO_3^-} + k_{\mathrm{NO}_x}}\right] \tag{22}$$

The uptake of each DIN species by each phytoplankton type is weighted by the concentration of each substrate. The inhibition of $NO_2^-$ and $NO_3^-$ assimilation in the presence of $NH_4^+$ had a negligible effect on the solutions and so was not included. Values for the maximum growth rate and the half-saturation constants were computed as functions of cell size following data-based allometric relationships in Litchman et al. (2007) as in Ward et al. (2012). The effective half-saturation constants for DIN uptake with respect to $\mu_{max}$ were calculated with respect to maximum uptake rate $V_{max}$ and minimum cell quota $Q_{min}$ from the relationships in Litchman et al. (2007), following Verdy et al. (2009) and Ward et al. (2012) (Table 2).

Light limitation was parameterized using an exponential form as a function of an instantaneous photosynthetic rate and the Chl *a* to Carbon ratio $\theta$, following Geider et al. (1997) and Hickman et al. (2010):

$$\gamma_{I_i} = 1 - \exp\left(\frac{-\Gamma_i \theta_i}{\mu_{max P_i} \gamma_{N_i} \gamma_T}\right) \tag{23}$$

Photosynthetic rate $\Gamma_i$ was computed as a function of photosynthetically active radiation $I(z)$, the maximum quantum yield of carbon fixation $\phi$ (mol C mol$^{-1}$ photons), and the absorption of light by phytoplankton $a_{P_i}^{\mathrm{chl}}$ (m$^2$ (mgChl)$^{-1}$) representing a mean value over all wavelengths, as:

$$\Gamma_i = \phi a_{P_i}^{\mathrm{chl}} I(z) \tag{24}$$

The Chl:C ($\theta$) varies with photoacclimation, and is computed using a steady-state solution Geider et al. (1997) as:

$$\theta_i = \frac{\theta_{max}}{1 + \frac{\Gamma_i \theta_{max}}{2(\mu_{max P_i} \gamma_{N_i} \gamma_T)}} \tag{25}$$

where $\theta_{max}$ is a maximum ratio.

**Loss rate** For the functional type populations represented with the Droop model (phytoplankton and heterotrophs), loss rate $L_i$ for type $i$ is parameterized with a quadratic mortality term, which represents mortality due to grazing and viral lysis. For mass balance, a specific mortality rate was calculated using

N-based biomass as a baseline, and so the specific loss rate for all biomasses $B_{i,j}$ and cell abundance $X_i$ of type $i$ is:

$$L_i = m_q B_{i,N} \gamma_T \tag{26}$$

Solutions were only slightly different in magnitude when choosing one of the other biomasses as the baseline (i.e. $m_q B_C$).

**Light** Light energy $I$ decreases with depth according to the attenuation coefficients for water $k_w$:

$$I(z) = I_{in} e^{(-z k_w)} \tag{27}$$

**Temperature** All microbial growth, grazing, and mortality rates are represented as a function of temperature (non-dimensional $\gamma_T$) using a formulation that follows the Arrhenius equation Dutkiewicz et al. (2015) as:

$$\gamma_T = \tau \exp(A_E(\frac{1}{T} - \frac{1}{T_0})) \tag{28}$$

where $T$ is the ambient temperature (K), $T_0$ is a reference temperature, $A_E$ regulates the temperature modification, and $\tau$ normalizes the maximum value. The model assumes a constant temperature profile, an average of the 10°S Pacific Ocean transect from the WOA 2013 climatology. This temperature dependency increases microbial rates by a factor of three from the deep to the surface, but does not impact solutions qualitatively.

**Vertical diffusion** A mixed layer was imposed by varying the vertical diffusion coefficient $\kappa_Z$ with depth, from a maximum $\kappa_{Zmax}$ at the surface to a minimum $\kappa_{Zmin}$, representing deep vertical mixing, with a length scale of $z_{ML}$. The conservation of total elements over the domain results in steady state accumulation of $POM$ at the bottom of the 2000m domain, conceptually representing a sediment layer. To avoid numerical instability, vertical mixing was increased within a bottom boundary mixed layer (100m length scale). The resulting vertical profile of $\kappa_Z$ ($m^2$ $s^{-1}$) is:

$$\kappa_Z = \kappa_{Zmax} e^{-\frac{z}{z_{ML}}} + \kappa_{Zmin} + \kappa_{Zmax} e^{-\frac{z-H}{100}} \tag{29}$$

where $z$ is in meters and H is the height of the domain (2000m).

**Nitrifying microorganisms** The parameterization of the nitrifying microorganisms was from Zakem et al. (2018). We did not resolve the quotas of these populations, and instead assumed that the C:N and C:P of nitrifier biomass is constant at $R_{bio_{CN}}$ and $R_{bio_{CP}}$, respectively. Because the biomass of these populations is relatively small, they have no impact on the solutions. A model run with only one DIN species and no nitrifiers resulted in nearly identical (visually indistinguishable) solutions.

**Numerical solution** Equations were integrated forward in time using a 4th order Runge-Kutta. Advection was carried out using the QUICK advection scheme, consisting of a linear interpolation between points weighted by an upstream 2nd order curvature, resulting in 3rd order accuracy. Fluxes were calculated at the faces of each grid cell, and concentrations at the centers.

### Text S3: Approximating steady state biomass

We first approximate a general expression for the steady state concentration of biomass. We combine Eqns. 1 and 2 as approximately $\frac{\partial B}{\partial t} = yVB - LB$, where yield $y \approx y_{maxC}$. We then exploit our use of a quadratic mortality term to describe loss rate $L$ as $m_q B$, which implicitly accounts for top-down control Steele and Henderson (1992); Edwards and Yool (2000), and solve for steady state biomass $B^*$ (µM N) as:

$$B^* = yVm_q^{-1} \tag{30}$$

We then define two types of limitations on bacterial biomass: by substrate concentration as "concentration-limited", and by consumption rate as "$V_{max}$-limited." For depleted substrates, we can assume $[DOC] << k_S$ and thus $V \approx V_{max} k_{DOC}^{-1} [DOC]$. This concentration-limited biomass can be approximated as:

$$B^*_{Conc} = yV_{max_{sp}} k_{DOC}^{-1} m_q^{-1} [DOC] \tag{31}$$

and so the amount of biomass sustained on a substrate is proportional to the concentration of that substrate. We note that because of the quadratic mortality assumption, this expression takes the relationship between substrate concentration and consumption rate one step further than $R^*$ and directly relate substrate concentration to biomass. As a result, a different parameterization of top-down control could decouple this relationship.

In contrast, for the biomass clades sustained on the recalcitrant pools of DOM, DOM concentrations are high, and so the consumption rate is saturated ($V \approx V_{max}$). The amount of biomass sustained on each recalcitrant DOM class is thus limited by the rate at which cells can process the substrate. This $V_{max}$-limited biomass can be approximated as:

$$B^*_{V_{max}} = yV_{max_{sp}}m_q^{-1} \quad (32)$$

### Text S4: Estimate of the results of Letscher and Moore (2015)

The gray shaded areas in Fig. 5 in the main text represent our estimate of the envelope of the C:N of remineralization of non-recalcitrant substrates and the magnitude of preferential remineralization by Letscher and Moore (2015). We use a tanh functional form to visually fit the results against an axis of neutral density, and then transform to depth coordinates using published code relating neutral density surfaces to depth (Jackett and McDougall 2002). For the calculation, we assumed a typical temperature profile for a stratified water column in the oligotrophic gyre from a cruise in the N. Pacific (Supplementary Figure 9 in Zakem et al. (2018)) and associated latitude and longitude.

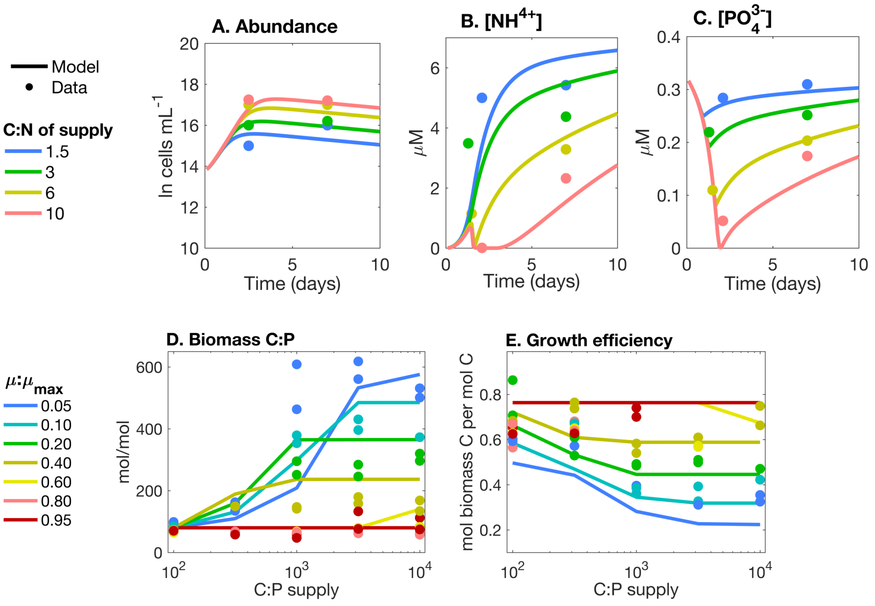


**Figure 1:** Metabolic model (lines) and observations (dots) from two experimental datasets: Goldman et al. (1987) (A-C) andGodwin et al. (2017) (D-E).

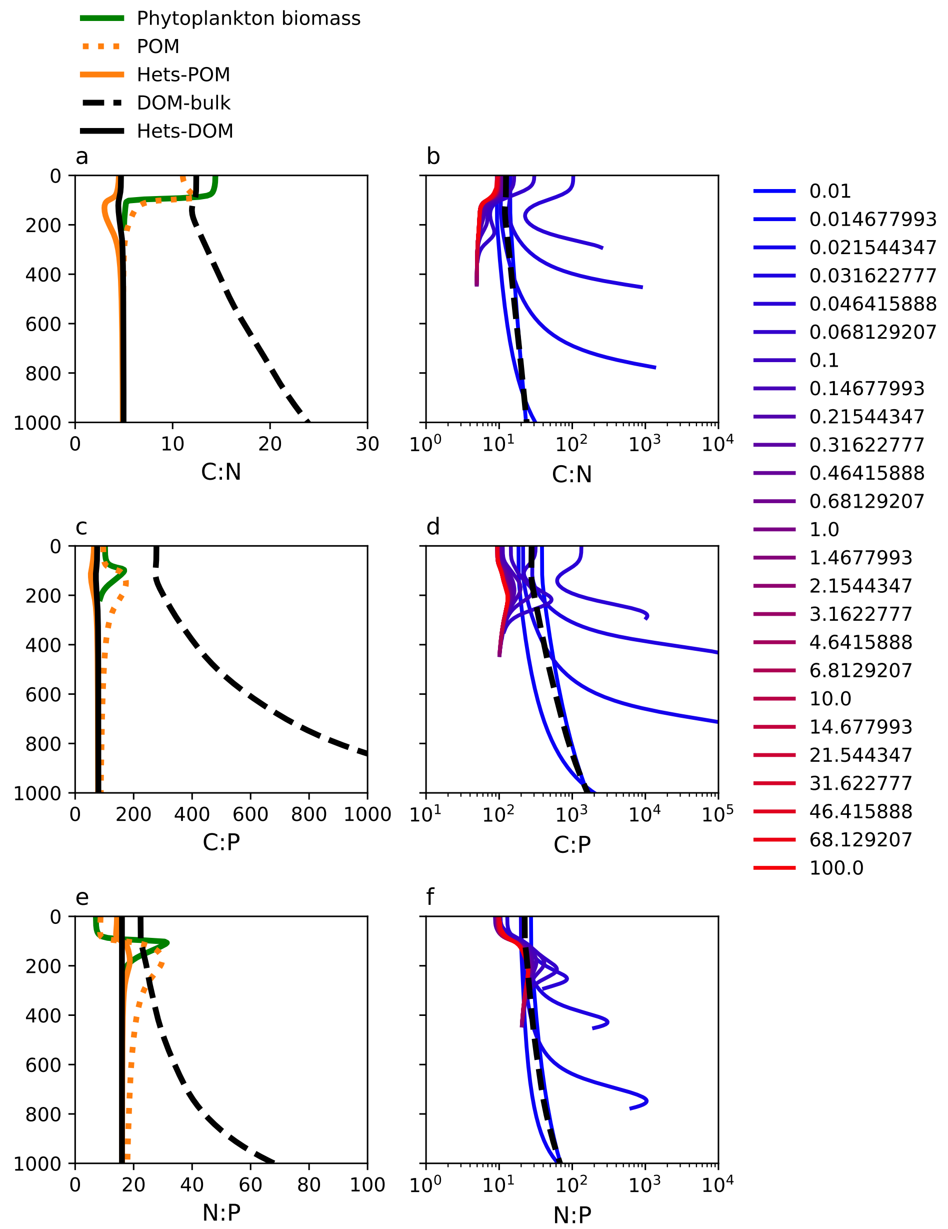


**Figure 2: All elemental ratios of the solutions.** The C:N, C:P, and N:P of biomass, POM, and bulk DOM are shown in the left column. Elemental ratios for each of the 25 DOM classes are shown in the right column. Primary production is N-limited in the model. Though, since DIC concentrations proportionally exceed DIP concentrations in the surface, the C:P of phytoplankton was also somewhat enhanced, allowing for the preferential remineralization of P relative to C for accumulated DOM classes. Solutions from Model C are illustrated.

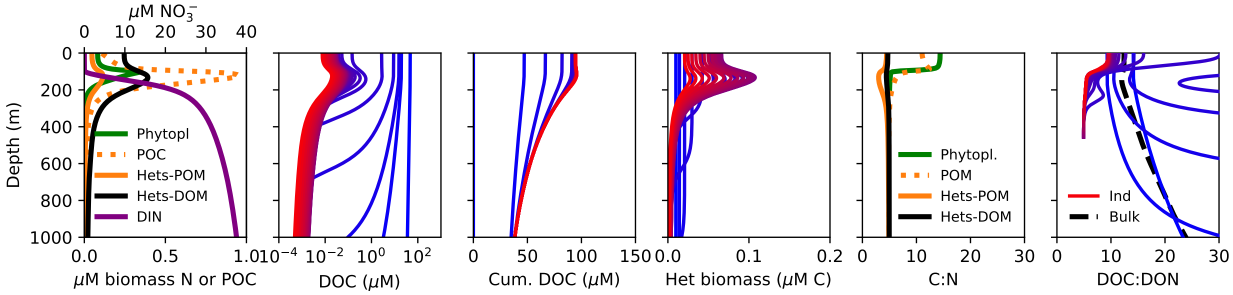


**Figure 3: Model C 'Default.'** For easiest comparison with the below sensitivity runs, we here illustrate the solutions from Model C in the main text.

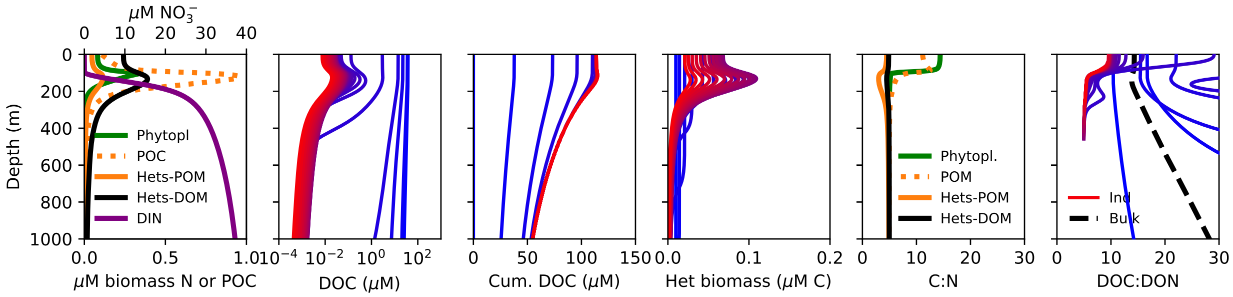


**Figure 4: No heterotrophic DIN uptake.** Heterotrophic populations are not allowed to assimilate DIN or DIP. These results are also illustrated in Fig. 1 in the main text. See also Fig. 14 for further model comparison.

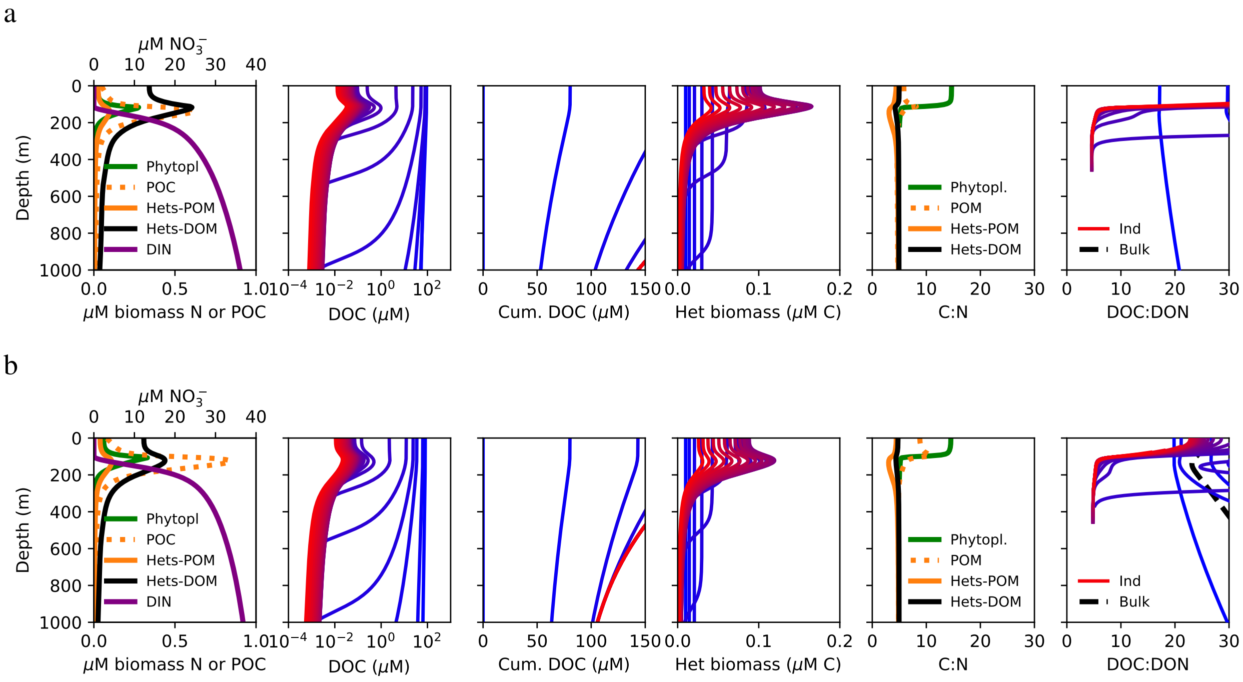


**Figure 5: Phytoplankton excretion.** For phytoplankton, the 'leakiness' parameter governing excretion $\gamma$ is 0.5 $d^{-1}$ in row a, and 0.1 $d^{-1}$ in row b.

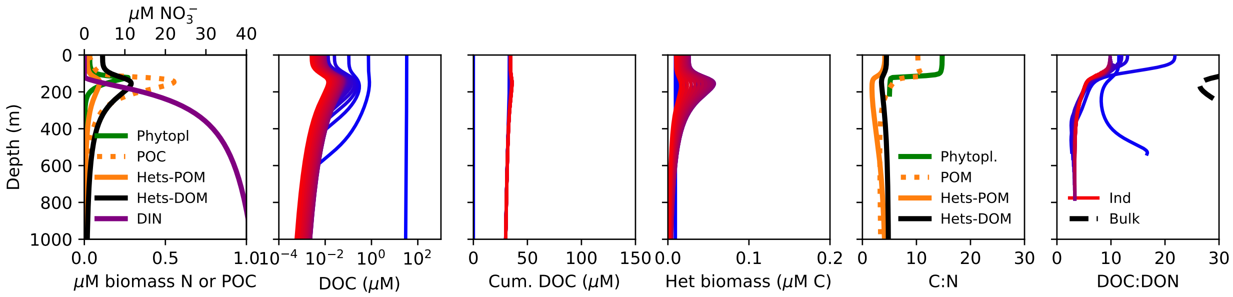


**Figure 6: Lower heterotrophic excretion.** For heterotrophs, the 'leakiness' parameter governing excretion $\gamma$ is 0.1 $d^{-1}$ rather than 0.5 $d^{-1}$ for the heterotrophic populations. Heterotrophic biomass C:N decreases to about 4.5 for the DOM-consumers and to about 4 for the POM-consumers.

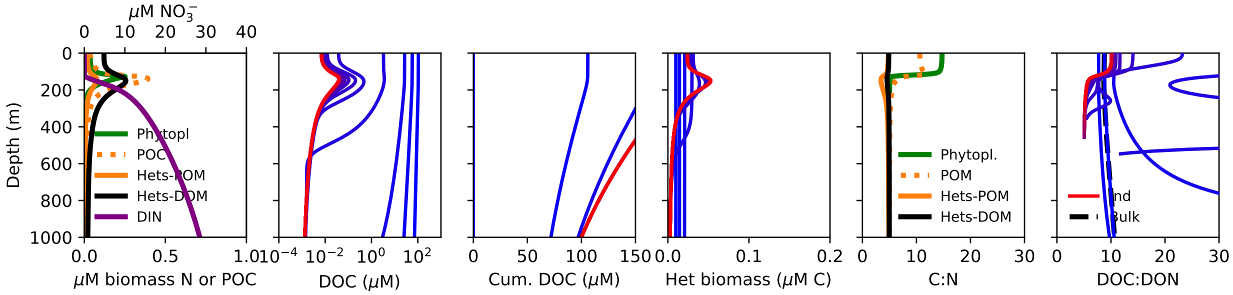


**Figure 7: Even distribution of DOM production.** The distribution of DOM production among the 25 classes is even, rather than lognormal.

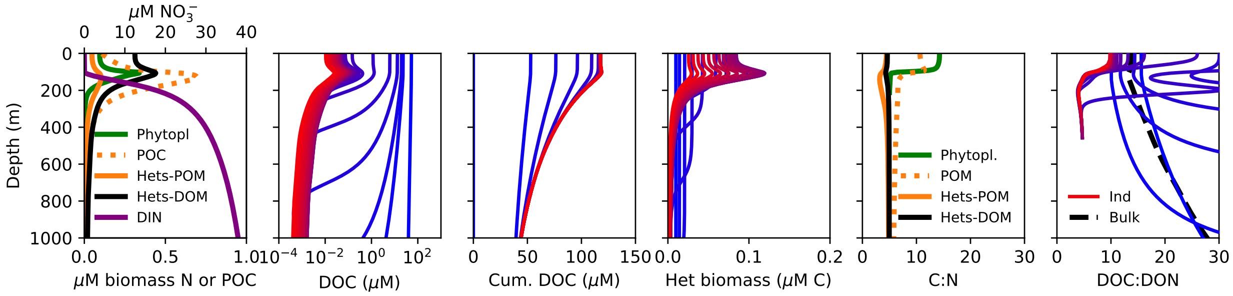


**Figure 8: DOM produced directly from mortality of biomasses.** Mortality from all biomass populations is partitioned equally into DOM and POM ($f_{mort} = 0.5$). Unlike the default model, no DOM is produced from the 'sloppy consumption' of POM. (Changing both of these at the same time keeps total DOM production similar, and allows for the largest contrast between the solutions.)

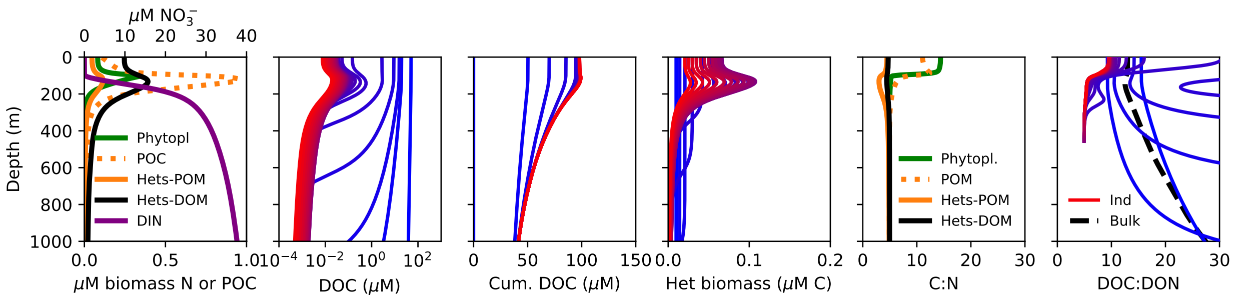


**Figure 9: Higher $Q_{min}$.** The minimum quota $Q_{minN}$ is 1 fmol N cell$^{-1}$, rather than 0.2 fmol N cell$^{-1}$.

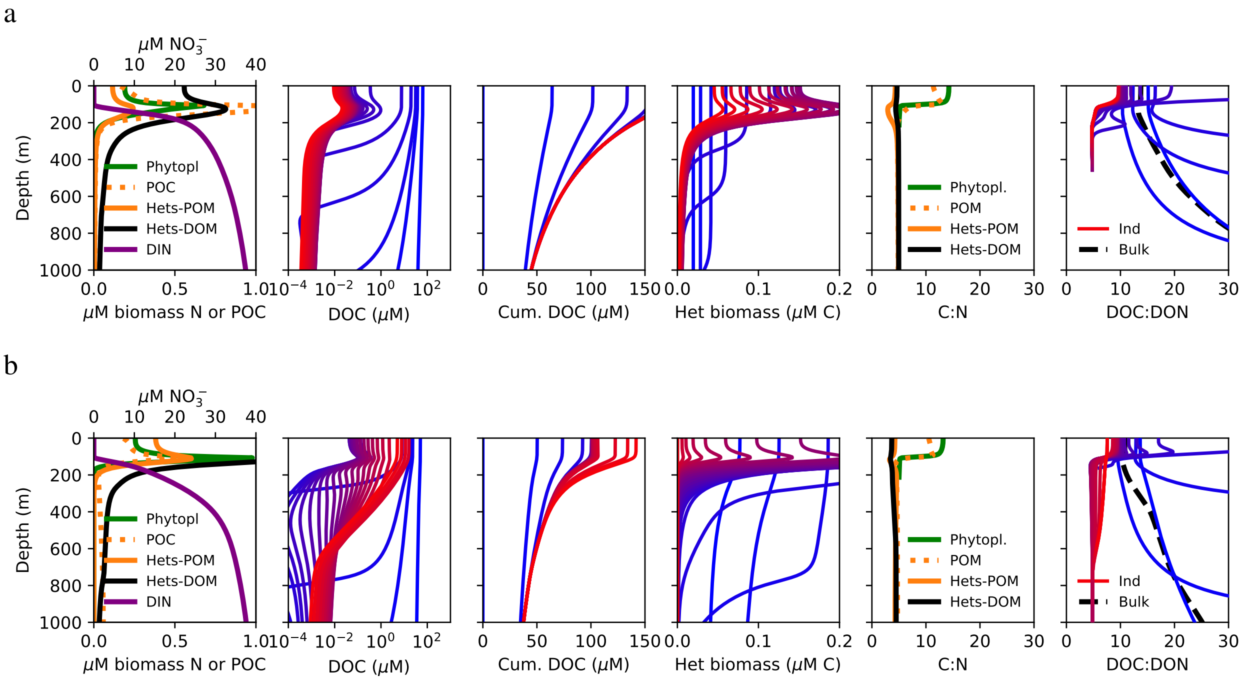


**Figure 10: Variation in mortality rate.** In row a, the quadratic mortality rate was reduced to half of its value (lower mortality rate). In row b, a linear mortality term is added to the biomass losses as 10% of the maximum growth rate for each population (increased mortality rate).

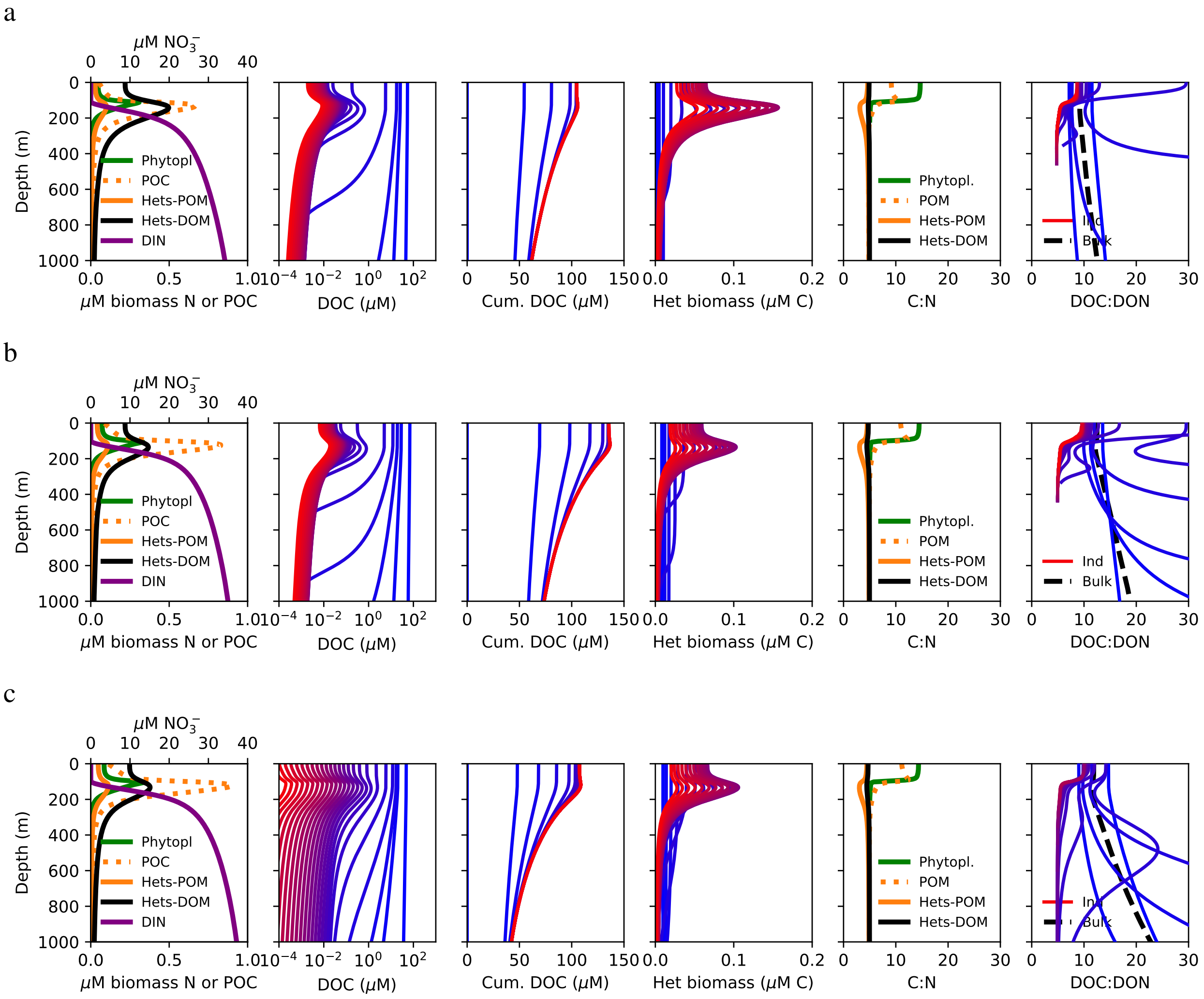


**Figure 11: Variation in heterotrophic growth rate parameters.** In row a, the maximum carbon yield is varied: $y_{maxC}$ increases linearly with increasing $V_{max}$, rather than being set at a constant value of 0.2 mol biomass per mol DOC consumed. For DOM class $i$ listed from slowest to fastest out of 25 total, $y_{maxC} = i/30$, resulting in values ranging from $y_{maxC} = 0.0333$ for the slowest DOM class to $y_{maxC} = 0.8333$ for the fastest DOM class. In row b, the maximum growth rate is varied: for each DOM-consuming population, $\mu_{max} = V_{max_{sp}}$. In row c, the half-saturation concentration is constant: for each DOM class, $k_C$ = 0.5 µM C, increasing the affinity of DOM with increasing uptake rate.

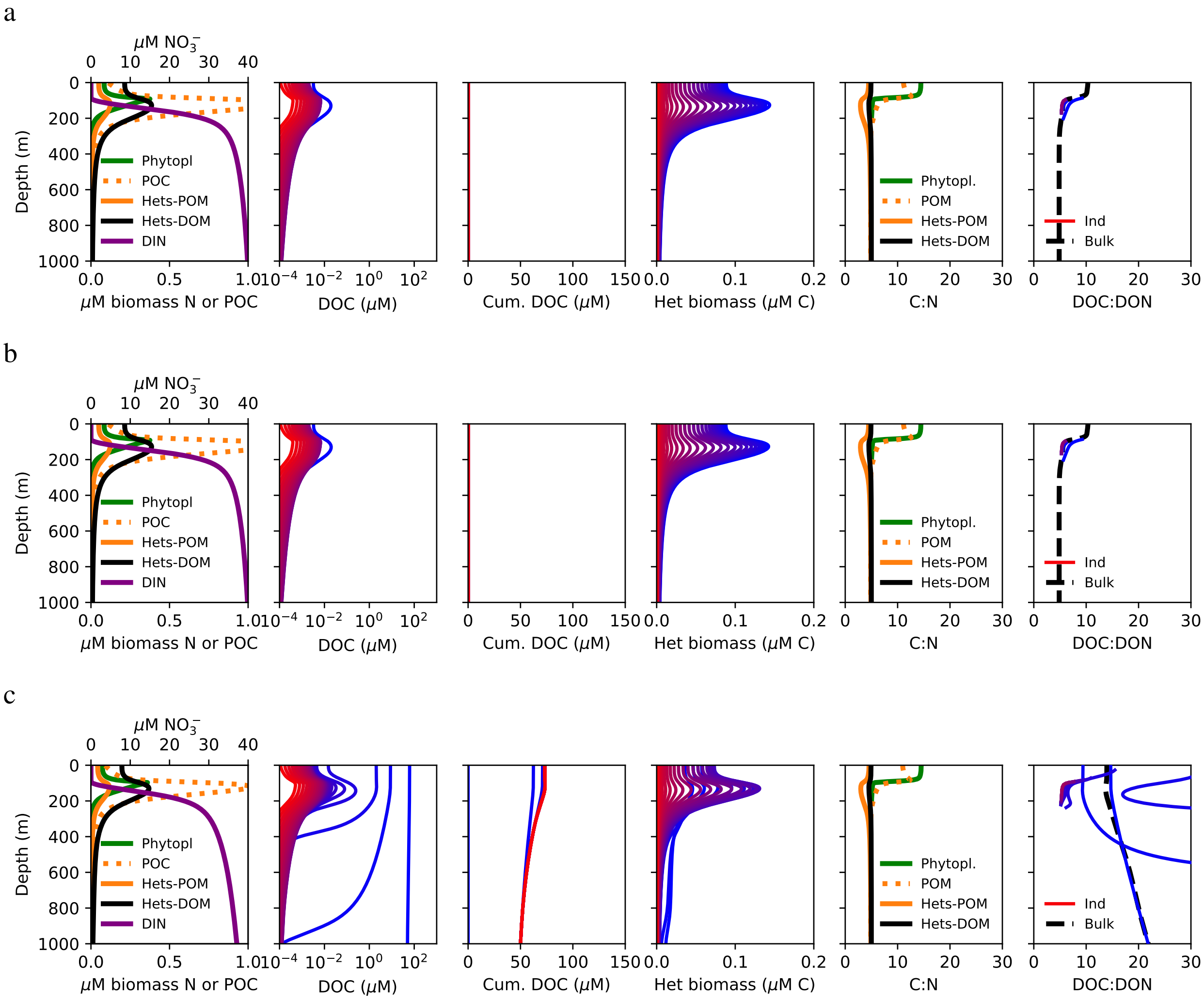


**Figure 12: Slow "generalist" to fast "specialist" configuration.** One of the 25 populations is able to consume all 25 DOM classes (blue line in biomass plot), another consumes the 24 most labile, another the 23 most labile, etc., so that one population (red line in biomass plot) consumes only the most labile DOM pool. Consumption remains at $V_{max_{sp}}$. In row a, $\mu_{max} = 1$ d$^{-1}$ for all 25 populations. In row b, $\mu_{max}$ for each population is the average of $V_{max_{sp_i}}$ for $i$ consumed substrates, and so $\mu_{max}$ increases with substrate specialization. In row c, $\mu_{max}$ equals the lowest value of $V_{max_{sp_i}}$ for a population taking up $i$ substrates, and so $\mu_{max}$ increases strongly with substrate specialization as a significant tradeoff.

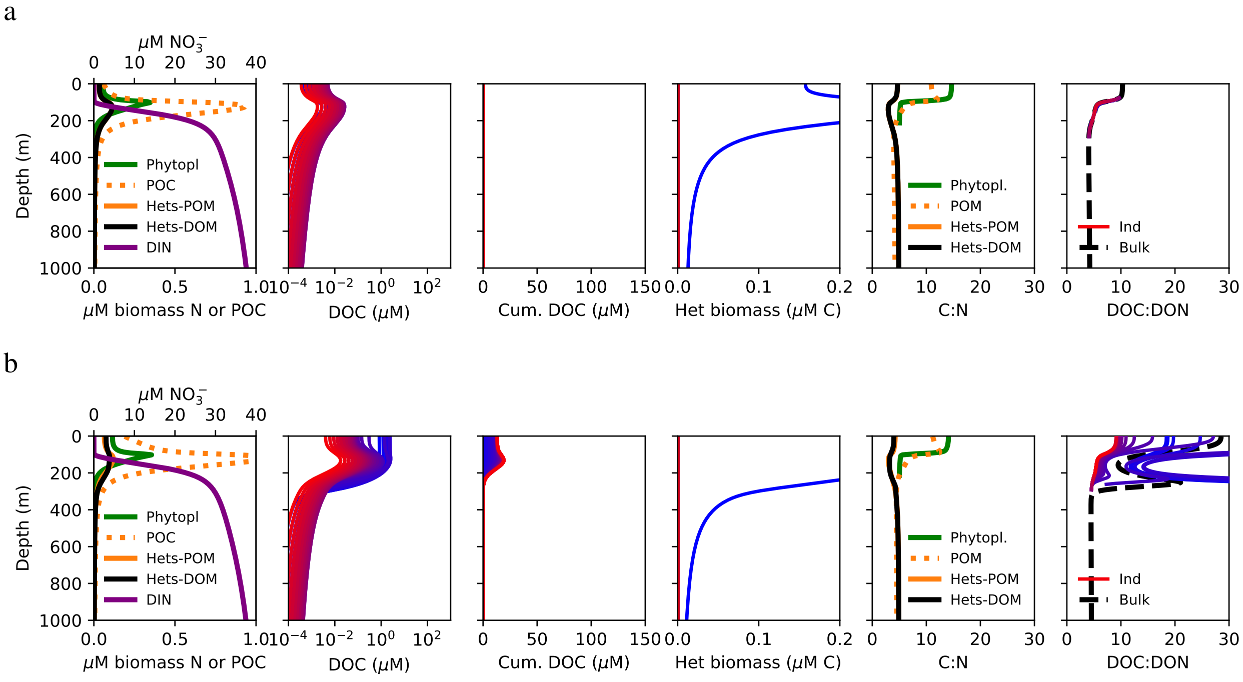


**Figure 13: One "generalist" DOM-consuming population.** In row a, $\mu_{max} = 1\ \mathrm{d}^{-1}$. In row b, $\mu_{max} = 0.1\ \mathrm{d}^{-1}$.

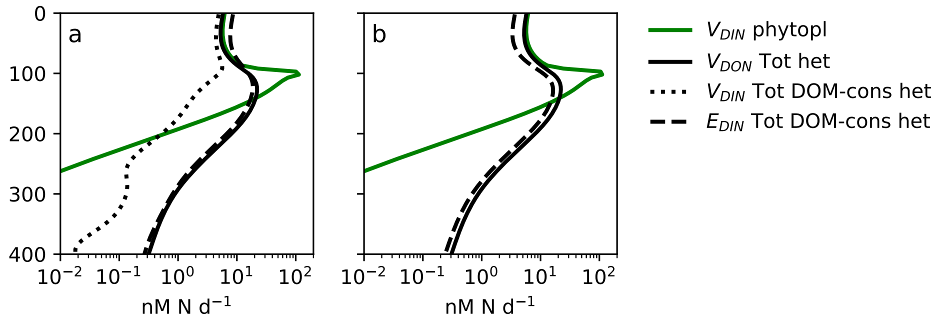


**Figure 14:** Comparison of N-cycling in the water column model with (a) and without (b) DIN assimilation by heterotrophic populations.

**Table 1:** Parameters for the case studies simulating two experimental datasets that differ from those of the water column model. (Otherwise, parameters are the same as in Table 1 and Table 2.)

| Parameter | Symbol | Value | Units |
|---|---|---|---|
| **Goldman et al. (1987):** | | | |
| Maximum specific uptake rate of $DOC_L$ | $V_{max_{spL}}$ | 10 | $d^{-1}$ |
| Maximum specific uptake rate of $DOC_S$ | $V_{max_{spS}}$ | 1 | $d^{-1}$ |
| Maximum growth rate | $\mu_{max}$ | 2 | $d^{-1}$ |
| Maximum carbon yield $DOC_L$ | $y_{maxC_L}$ | 0.6 | unitless |
| Maximum carbon yield $DOC_S$ | $y_{maxC_S}$ | 0.2 | unitless |
| Linear mortality rate | $m_l$ | 0.1 | $d^{-1}$ |
| **Godwin et al. (2017):** | | | |
| Maximum specific uptake rate of DOC | $V_{max_{sp}}$ | 10 | $d^{-1}$ |
| Maximum growth rate | $\mu_{max}$ | 3.9 | $d^{-1}$ |
| Maximum carbon yield | $y_{maxC}$ | 0.8 | unitless |

**Table 2:** Parameters for the water column model simulation illustrated in the main text. The variations in parameter values that are illustrated in Supplement 4 are indicated in parentheses.

| Parameter | Symbol | Value | Units |
|---|---|---|---|
| **DOM uptake:** | | | |
| Maximum specific uptake rate (Fig. 3) | $V_{max_{sp}}$ | 0.01 – 100 | $d^{-1}$ |
| DOC half-saturation | $k_C$ | $0.5V_{max_{sp}}$ | µM |
| **POM uptake:** | | | |
| Maximum specific uptake rate | $V_{max_{sp}POM}$ | 1 | $d^{-1}$ |
| POC half-saturation | $k_{POC}$ | 0.5 | µM |
| Production of DOM from POM 'sloppy feeding' | $\alpha$ | 2 | mol DOM mol $POM^{-1}$ |
| **Inorganic uptake:**§ | | | |
| Maximum specific DIN uptake rate | $V_{max_{sp}DIN}$ | 8.8 | $d^{-1}$ |
| $NO_x^-$ half-saturation | $k_{NO_x}$ | 81 | nM |
| $NH_4^+$ half-saturation | $k_{NH_4}$ | 41 | nM |
| Maximum DIP uptake rate | $V_{maxDIP}$ | $R^{-1}_{bio_{NP}} V_{maxDIN}$ | $d^{-1}$ |
| $PO_4^{2-}$ half-saturation | $k_{PO_4}$ | $R^{-1}_{bio_{NP}} k_{NH_4}$ | nM |
| **Heterotrophic growth:** | | | |
| Maximum growth rate | $\mu_{max}$ | max(1,$V_{max_{sp}}$) | $d^{-1}$ |
| Maximum carbon yield (water column) | $y_{maxC}$ | 0.2 (0.0333–0.8333) | unitless |
| Regulation rate of excretion | $\gamma$ | 0.5 (0.1) | $d^{-1}$ |
| **Phytoplankton growth:** | | | |
| Maximum growth rate | $\mu_{maxP}$ | 2 | $d^{-1}$ |
| Maximum DIC uptake rate | $V_{maxDIC}$ | $R_{bio_{CN}} V_{maxDIN}$ | $d^{-1}$ |
| PAR half-saturation for autotrophy | $k_I$ | 10 | W $m^{-2}$ |
| Regulation rate of excretion | $\gamma$ | 0 (0.5) | $d^{-1}$ |
| **Grazing and mortality:** | | | |
| Quadratic mortality rate | $m_q$ | 1 (0.5) | µM $N^{-1}$ $d^{-1}$ |
| Fraction of mortality to DOM vs. POM | $f_{mort}$ | 0 (0.5) | unitless |
| **Nitrifying microorganism growth:**† | | | |
| $NH_4^+$ yield, AOO | $y_{NH_4}$ | $112^{-1}$ | unitless |
| $NO_2^-$ yield, NOO | $y_{NO_2}$ | $334^{-1}$ | unitless |
| Maximum $NH_4^+$ uptake rate, AOO | $V_{max NH_4,AOO}$ | 50.8 | mol $NH_4^+$ mol $N^{-1}$ $d^{-1}$ |
| Maximum $NO_2^-$ uptake rate, NOO | $V_{max NO_2,NOO}$ | 23.6 | mol $NO_2^-$ mol $N^{-1}$ $d^{-1}$ |
| $NH_4^+$ half-saturation, AOO | $K_{NH_4,AOO}$ | 133 | nM |
| $NO_2^-$ half-saturation, NOO | $K_{NO_2,NOO}$ | 287 | nM |
| **Temperature dependence:**‡ | | | |
| Reference temperature | $T_0$ | 293.15 | K |
| Temperature regulation | $A_E$ | -4000 | K |
| Temperature normalization | $\tau$ | 0.8 | unitless |
| **Physical parameters:** | | | |
| Maximum incoming PAR flux | $I_{max}$ | 1400 | W $m^{-2}$ |
| PAR attenuation in water | $k_w$ | 0.04 | $m^{-1}$ |
| Mixed-layer attenuation depth | $z_{ML}$ | 20 | m |
| Minimum vertical mixing coefficient | $\kappa_{Zmin}$ | $5 \cdot 10^{-5}$ | $m^2$ $s^{-1}$ |
| Maximum vertical mixing coefficient | $\kappa_{Zmax}$ | $10^{-2}$ | $m^2$ $s^{-1}$ |
| $POM$ sinking rate | $w_s$ | 10 | m $d^{-1}$ |

§Calculated for a 0.5 $\mu$m cell diameter following Litchman et al. (2007) and Ward et al. (2012).
†Zakem et al. (2018).
‡Dutkiewicz et al. (2015).
